# Validating the Nordic Thingy:53 as an Accessible Tool for Equine Behavioural Analysis using Edge AI

**DOI:** 10.64898/2025.12.05.692598

**Authors:** Rhune Vanden Eynde, Ivan Herrera Olivares, Lucinda Kirkpatrick

## Abstract

Understanding animal movement and behaviour is integral to ensuring welfare needs are met, a central goal within the scientific community. Among domesticated animals, equids are unique; they serve as both companion and production animals, with estimates showing around 112 million working equids supporting 600 million people worldwide. Furthermore, the horse sport industry is a global multi-billion-dollar sector. However, these roles inevitably expose horses to potential physical and psychological harm. Consequently, being able to monitor behaviour accurately is critical to meeting their welfare needs.

Here, we describe the design and evaluation of machine learning models capable of recognizing equine gaits and (semi-)natural behaviours using the Edge AI platform, Edge Impulse. Accelerometer motion data were collected using the Nordic Thingy:53 prototyping platform located on the chest of the horse. Both gaits and behaviours were detected with high mean training and testing accuracies (>95.00%), including idle, walk, trot, canter, fly-twitch, eating hay, head down or up, grazing, and rolling. While transformation types (Fast Fourier Transform (FFT) vs. Wavelet transform) were compared, they were not found to have a drastic effect on model performance. Instead, model accuracy and sensitivity were most impacted by the volume of training data included. This suggests that while transformations may refine model selection, sufficient training data is the single most important factor in defining effective Edge AI models. This study extends the evidence detailing the utility of accelerometers for equine movement and demonstrates that the accessible Nordic Thingy:53 platform can be used to develop highly accurate models for behavioural analysis. This is an important first step towards building effective monitoring methods that can recognise subtle behavioural changes that indicate alterations to emotional or physical states in equines.

## 1. Introduction

In recent decades, the topic of animal welfare has received increasing interest from the scientific community (Khillare & Kaushal, 2021). Current knowledge demonstrates that simply ensuring the basic physiological needs of the animal are met does not guarantee positive animal welfare (Hewson, 2003; Hockenhull & Whay, 2014; Whay et al., 2003). Consequently, modern welfare assessment frameworks have evolved to incorporate the animal’s emotional state. Since emotional states cannot be measured directly, they must be inferred through measurable proxies such as the animal’s ability to perform its natural behaviour repertoire or freedom from pain induced movement alterations (Khillare & Kaushal, 2021; Webster, 2016).

Healthy, stress-free animals typically divide their time between activities that satisfy basic requirements for movement, food and rest, usually exhibiting highly repetitive daily routines (Auer et al., 2021). Therefore, time budget analyses, which classify behavioural activities over time, have become a cornerstone of welfare assessments (Aoun et al., 2024; Auer et al., 2021; Flannigan & Stookey, 2002; Huettner et al., 2021; Yarnell et al., 2015). Historically, these studies relied on direct animal observation (Huettner et al., 2021; Mench & Mason, 1997; Tadich et al., 2013; Zupan et al., 2020), a method that is labour intensive, limited to daylight hours and prone to observer bias (Eerdekens et al., 2020).

To overcome these limitations, researchers are increasingly adopting artificial intelligence (AI) derived approaches (Ezanno et al., 2021). While AI monitoring is well established in wildlife research (Ladds et al., 2018; Nathan et al., 2012; Shepard et al., 2008), its application to equine welfare remains underutilised compared to other domestic species (Hockenhull & Whay, 2014, Zhang et al., 2023). Negative welfare in horses is often indicated by a shift away from natural behaviour patterns, such as reduced grazing time or altered sleep intervals, or the emergence of abnormal behaviours (Auer et al., 2021; Benhajali et al., 2007). Capturing these subtle shifts requires detailed surveillance over 24 hour cycles, a task for which biotelemetry and AI are uniquely suited, offering higher accuracy and temporal resolution than human observers can, at considerably lower economic costs (Eerdekens et al., 2020; Ladds et al., 2018).

However, a barrier to the widespread adoption of these technologies is often the cost and technical complexity of the equipment. Previous studies have shown promise using AI for grazing (Maisonpierre et al., 2019; Weinert et al., 2020) and gait determination (Eerdekens et al., 2020) but there is a need to validate accessible, user friendly platforms that can bring these capabilities to wider welfare applications.

In this study, we aim to address this gap by evaluating the Nordic Thingy:53 prototyping platform (hereafter referred to as Thingy:53; https://www.nordicsemi.com/Products/Development-hardware/Nordic-Thingy-53), a rapid prototyping platform combined with Edge Impulse (https://edgeimpulse.com/), a user friendly machine learning interface. Our goal was to produce models capable of reliably determining the gaits and natural behaviours of domesticated horses (*Equus ferus caballus*) to support objective welfare assessments. We utilised a three step approach to refine our models:

- Gait detection, training a model to accurately identify gaits (walk, trot, canter) during controlled lunging,
- Guided behaviour: Training a model to detect specific guided movements (e.g. head movements),
- Natural behaviour: validating the model’s ability to detect spontaneous natural movements (e.g. grazing, resting, rolling) while the horse was at pasture.

## 2. Materials and methods

### 2.1 Data collection

Equine motion data was collected at the Flaming Star Ranch (Hamont-Achel, Belgium). The Nordic Thingy:53, a compact, multi-sensor prototyping platform was housed in a custom harness secured to the horse’s chest, positioned centrally above the pectoral muscles (Supplementary Data, Figure S1). This location was chosen to minimise oscillation while remaining non-invasive. Data were transmitted via Bluetooth Low Energy (BLE) to the nRF Edge Impulse mobile application and uploaded to the Edge Impulse Studio for model training.

### 2.2 Experimental protocols

#### 2.2.1 Gait determination (lunging)

Data were collected from 5 horses. Horses were lunged in circles (left and right reins) at four gaits: idle, walk, trot and canter. A sampling frequency of 20kHz. Due to varying cooperation levels, the duration of data collection varied per horse (Supplementary data: Table S1).

#### 2.2.2 Controlled behaviour

To validate the detection of specific activities, five horses were guided into performing discrete behaviours. Head movements (up / down / left / right) were induced using food rewards, mimicking movements seen in the field. Specific leg movements were captured by leading, and “eating hay” was initiated by providing horses with hay in their stalls, either on the ground or in a haynet. Grazing in this specific experiment included both the act of cropping grass and associated limb movements.

#### 2.2.3 Natural behaviour

To validate the system in a real word setting, three horses were monitored in individual pastures between 09:00 and 17:00 (August-November 2023). Horses had ad libitum access to water and were provided with hay and feed twice a day. Natural behaviours were observed and collected in one minute intervals using the Thingy:53 couple with the nRF Edge Impulse mobile application, coupled with voice notes detailing the observed behaviours. Samples were then manually split into different behaviours for analysis using the split-sample function in Edge Impulse (Figure S3, Supplementary data). In this experiment, grazing was strictly defined as the act of cropping / chewing grass. To ensure data quality, behaviour classes with insufficient instances / not observed across multiple subjects were excluded from the final analysis.

### 2.3 Model construction and refinement

Machine learning models were developed within the Edge Impulse platform. To test the hypothesis that this technology would be accessible to non experts, we prioritised a “low adjustment” approach, retaining factory settings where possible. Adjustments were limited to:

1. Spectral analysis type: Comparison between Fast Fourier Transform (FFT) and Wavelet transform
2. Window size: The extent of the data to be processed per classification, in milliseconds. Stepwise optimization was carried out based on training accuracy (Supplementary data, Figure S4, Tables S4-S5) controlled and natural behaviour. For gait determination models, window size was fixed at 1000ms (FFT) and 8500ms (Wavelet, corresponding to a completed lunging circle while cantering). Model training accuracy values were used to identify the optimal window size.

### 2.4 Model selection and metrics

Model performance can be evaluated using four key metrics: Accuracy, Precision, Recall and the F1 score. Model accuracy measures the number of times the model accurately predicts the outcome, providing values for both training and testing datasets (Sweeney et al., 2022). Comparing both values is critical; high training accuracy that does not mirror high testing accuracy would suggest overfitting; therefore an optimal model should demonstrate high accuracy for both model training and model testing (Ying, 2019). However, relying solely on accuracy can be misleading in behavioural studies when datasets are imbalanced, as it treats all classes as being equally important and represented. This does not occur when certain behaviours are rarer (e.g. horses will spend significantly more time grazing than rolling). In such cases a model could achieve high accuracy by simply ignoring rare behaviours (Sweeney et al., 2022).

To address this, we incorporated precision, a measure of the reliability of positive detections, and recall, ability to find all instances (Sweeney et al., 2022). Because no single metric is perfect, we also considered the F1 score for model selection. The F1 score computes the harmonic mean of precision and recall, providing a single metric that balances the trade off between missing a behaviour and misidentifying one (Sweeney et al., 2022). Given the unbalanced nature of the datasets in this study, we prioritised model accuracy and the F1 score together for model selection.

#### 2.4.1 Gait determination experiment

A two-stage model selection approach was adopted. First, the effect of including lunging direction information on model performance was evaluated. Second, stepwise increases in training data volume was carried out by including data from a second and third horse (Figure S4, Supplementary data). In total, 14 preliminary models were created and compared, 7 using FFT and 7 using Wavelet transforms (Figure S4, Table S10, supplementary data).

#### 2.4.2 Controlled and Natural behaviour experiment

After initial refinement, models were optimised by removing or combining variables in a stepwise fashion (Figure S5, Supplementary data). Selection criteria included: data availability, F1-score performance, similarity to other movements (confusion risk) and the number of subjects from which the movement was collected. This process yielded a refined candidate model set (“Model Set 2”) consisting of:

- Controlled behaviour: 7 FFT and 7 Wavelet models (supplementary data, Table S6)
- Natural behaviour: 22 FFT and 20 Wavelet models (supplementary data, Table S7)

### 2.5 Model testing and robustness

#### 2.5.1 Gait determination experiment

The robustness of the selected gait models was evaluated by testing against the inclusion of novel data (models 8-11) where subjects were shifted between training and test sets, and then to reduced novel data (model 12, horse with limited data samples in the test set). This was repeated for both transform types, resulting in 10 final models (5 FFT, 5 Wavelet) retaining all gait classes (Figure S4, Table S10, supplementary data).

#### 2.5.2 Controlled and Natural behaviour experiment

To rigorously test the robustness of the controlled and natural behaviour experiments to novel subjects, specific horses were shifted between the training and test datasets (“model set 3”, Figure S5, supplementary data). Data were initially collected from five horses for the controlled behaviour experiment, but only three individuals performed the complete repertoire of target movements. To ensure testing metrics represented the model’s ability to detect all behaviours, only the three individuals that were fully represented across all behavioural categories were shifted between training and testing datasets. For the natural behavioural experiments, sufficient data was present across the target behaviours to include all three horses (see Table S8 and S9 in supplementary data for an overview of the movements and behaviours in model set 3 alongside an overview of the data collected per behaviour and horse).

## 3 Results

All three experimental protocols yielded at least one model capable of reliably recognising the respective movements, defined as maintaining a mean training and testing accuracy value > 95.00%.

### 2.1 Gait determination experiment

The optimisation process yielded highly robust models for gait classification. Most candidate models exceeded 90.00% training accuracy and 80.00% testing accuracy (Figure 1, Table 1). Based on the combined metrics (Accuracy, F1-score and performance on novel data, model 11 (3 horse training set) was identified as the top FFT model, and model 8 (2 horse training set) as the top wavelet model. Both models exhibited distinct feature clustering (Figure S6, Supplementary data) and successfully classified all four gaits with mean accuracies exceeding 98.00% (Figure 2).

**Table 1:**
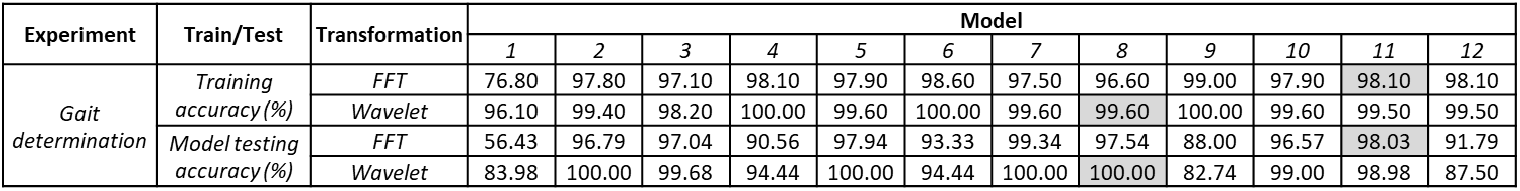
Overview of the training and model testing accuracy values obtained for the gait determination models. Accuracies of the top models are indicated in grey.

**Figure 1.**
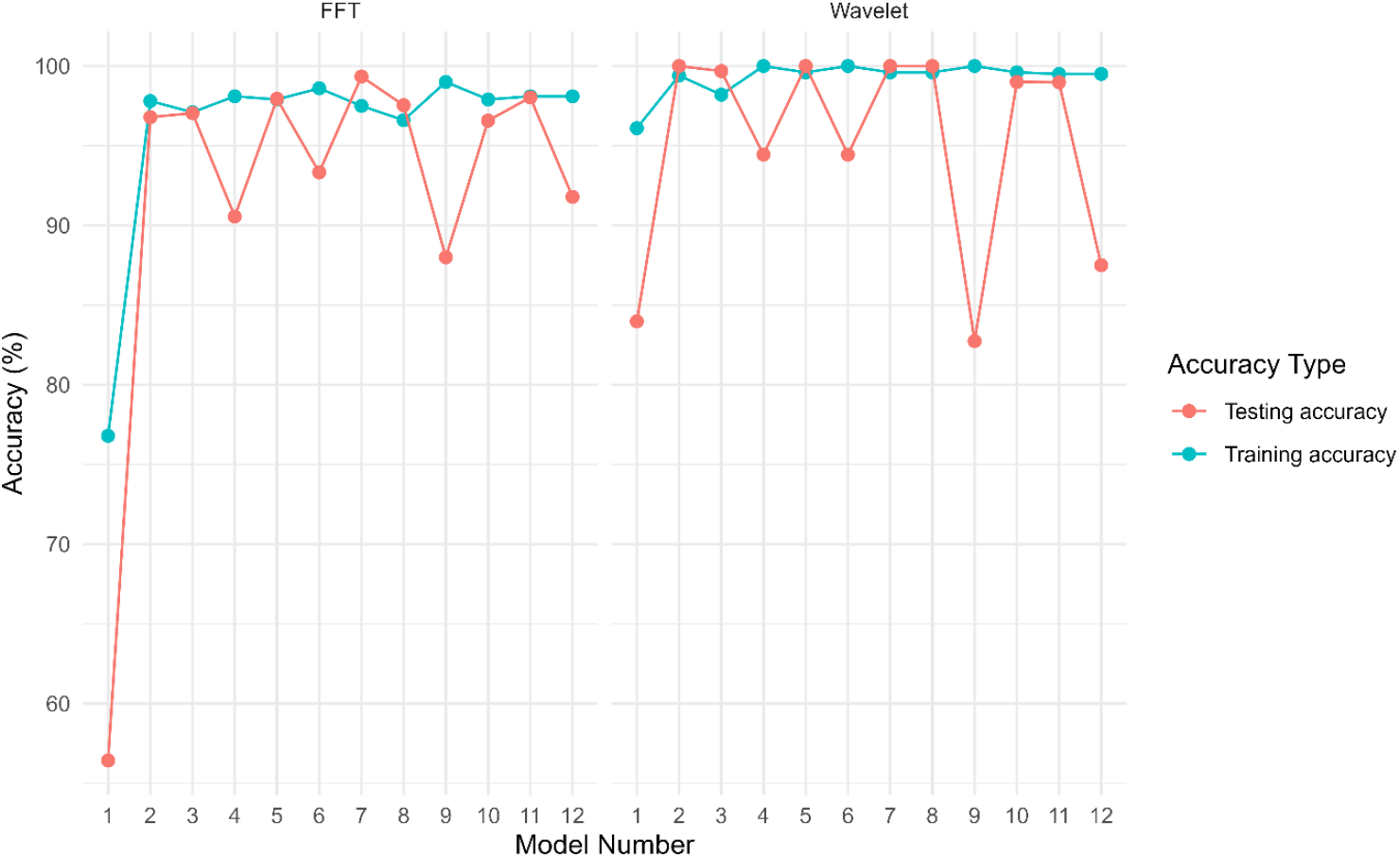
Overview of the model training accuracy values (indicated in blue) and model testing accuracy values (indicated in orange) obtained for the 24 gait determination models. Training accuracy values indicate how well the model is able to classify data contained in the training set. Model testing accuracies indicate how well the model is able to classify novel data contained in the test set. Accuracy values on the Y-axis start at 50.00%. A distinction is made based on transformation type, models containing a Fast Fourier transform are located on the left, models containing a Wavelet transform are located on the right.

**Figure 2.**
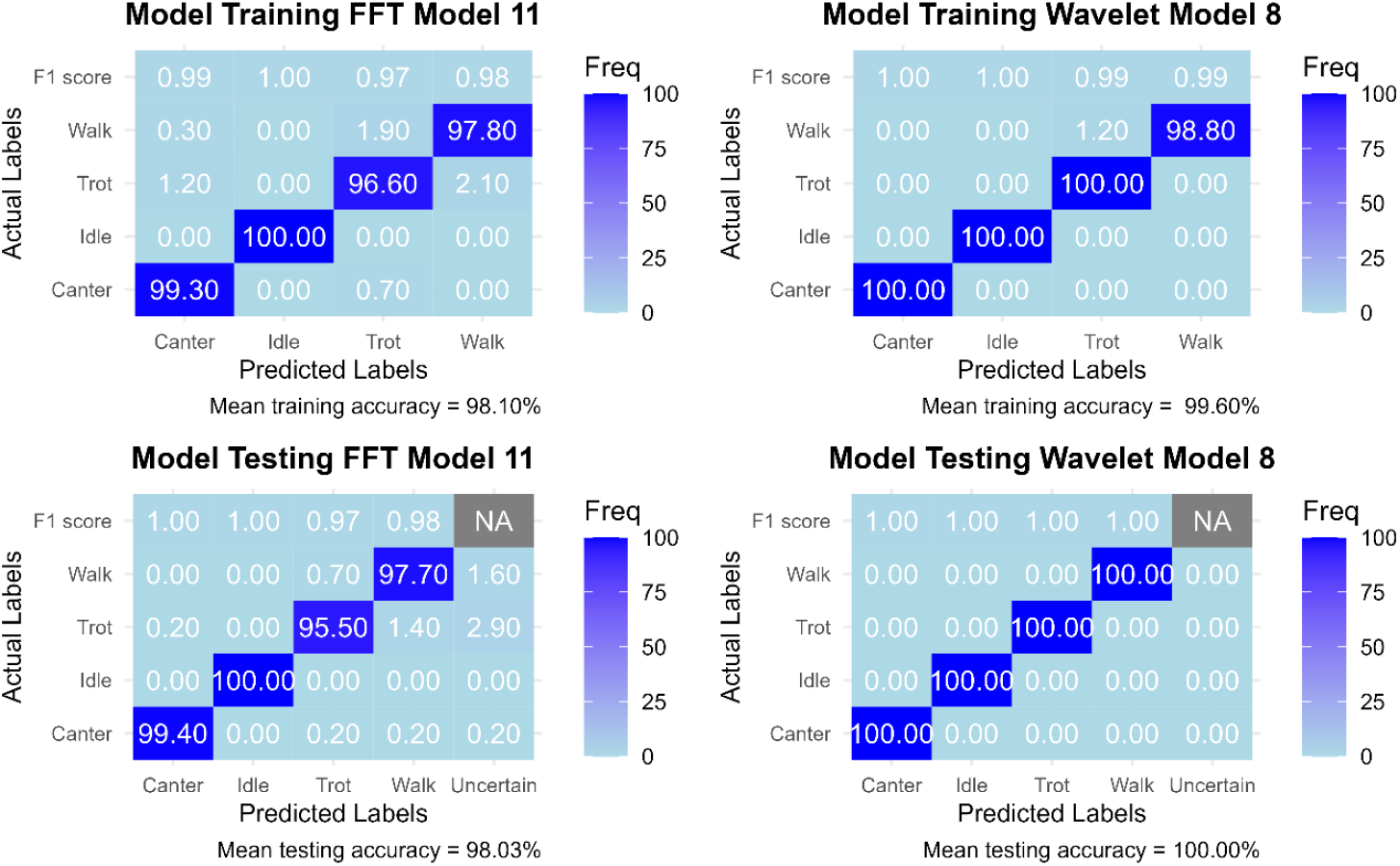
Overview of the confusion matrices pertaining to both the model training and testing accuracy values of gait determination FFT model 11 and Wavelet model 8. Accuracies are given in percentages, F1 scores are given as values between 0 and 1 with higher accuracies and F1 scores indicating better model performance. The colour-gradient shown on the frequence bar indicates the classification percentage per square, with darker blue indicating a larger number of observations being classified as the corresponding movement/behaviour. Correct classifications are located on the diagonal. Mean training and testing accuracies per model are also depicted below the matrices.

### 3.2 Controlled and natural behaviour experiments

#### Model refinement

Window size optimisation resulted in an 18,000ms window for controlled behaviour for both FFT and Wavelet transformed models (Table S4, supplementary data), while for natural data a window size of 4000ms was selected for FFT models and 3800ms for Wavelet transformed models (Table S5, Supplementary data).

#### Model selection

Stepwise refinement of controlled behaviour models increased training accuracy from ∼90.00% to ∼99.00%, with model 7 selected as the final candidate for both FFT and Wavelet transforms, achieving perfect F1 scores on the training data (1.0, Figures S7-S8, Table S8, Supplementary data). Within the natural behaviour experiment, training accuracy improved from ∼80.00% to ∼93.00%. Model 22 was selected as the final candidate mode, achieving F1-scores ranging from 0.77-1.0 across classes (Figure 3, Figures S9-S10, Table S7, Supplementary data).

**Figure 3.**
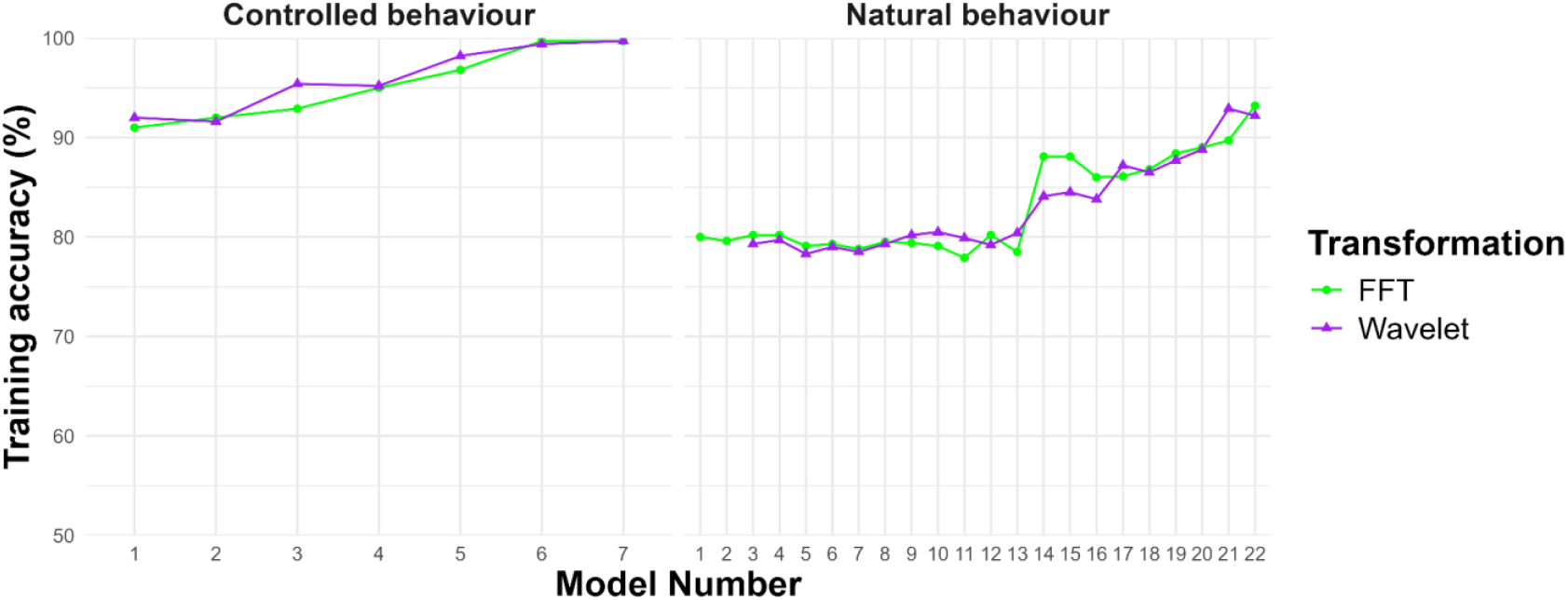
Overview of the training accuracy values obtained during model selection for the controlled behaviour experiment (indicated in blue) and natural behaviour experiment (indicated in red). Training accuracy values on the Y-axis start at 50.00%. A distinction is made between models using an FFT and models using a Wavelet transform.

#### Model testing for robustness to novel data

Final models were subjected to sensitivity to novel data testing (model set 3), where subjects were separated between training and testing sets. For the controlled behaviour experiment, while models classified known movements with >95.00% accuracy, performance dropped when applied to novel subjects and was also highly dependent on which novel subject was included / excluded and differed between transformation type (Figure 4; FFT model accuracy ranged between 99 – 82% test accuracy, Wavelet model ranged between 98 – 65% test accuracy, Table S8, S11, Supplementary data). Cluster analysis (Figures S11-12, Supplementary data) revealed that while distinct movements like “rolling” and “head up / down” were clearly separated, biologically similar movements such as “grazing” and “eating hay” were harder to distinguish.

**Figure 4.**
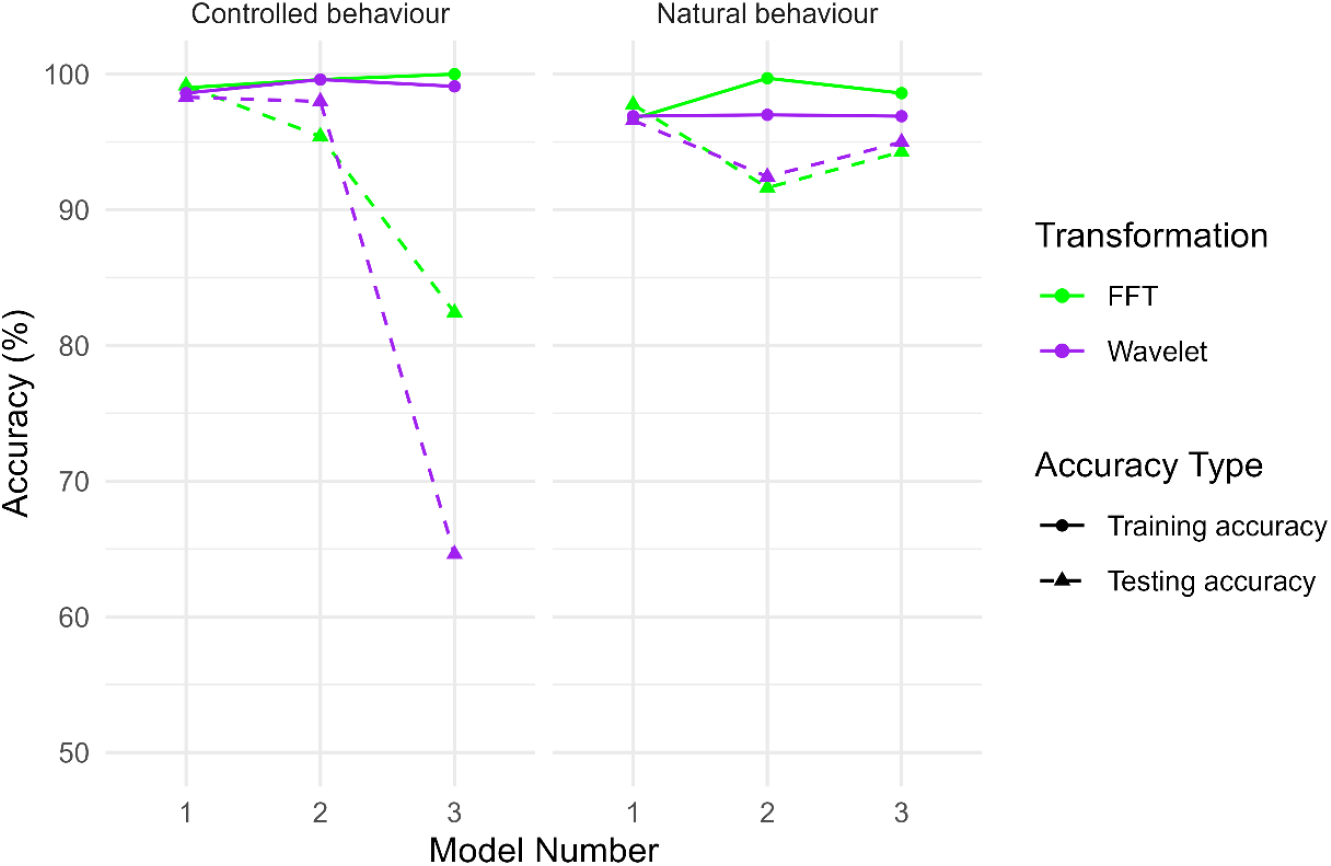
Overview of the training accuracy values (A) and the model testing accuracy values (B) obtained for model set 3 belonging to the controlled behaviour experiment (indicated in blue) and natural behaviour experiment (indicated in red). Accuracy values on the Y-axis start at 50.00%. A distinction is made between models using an FFT and models using a wavelet transformation.

The natural behaviour models demonstrated high robustness to novel subjects, correctly categorising events in >90.00% of cases and similar performances between Wavelet and FFT transformations (Figure 5). Classes “grazing” and “idle” were excluded from these analyses because they were not observed across all three test subjects (see Tables S9, S11 for overview of the horses, movements and data included, Figure S4). Cluster analyses showed good separation for the selected behaviours, with the top Wavelet model able to distinguish between the walking and eating hay movement and the head down/head up and flytwitch movement (see Figures S13 and S14, supplementary data, for more information). Both top models showed some overlap between the head down/head up movement, flytwitch and eating hay movement. However, the walking movement showed little to no overlap with the other three movements in either model.

**Figure 5.**
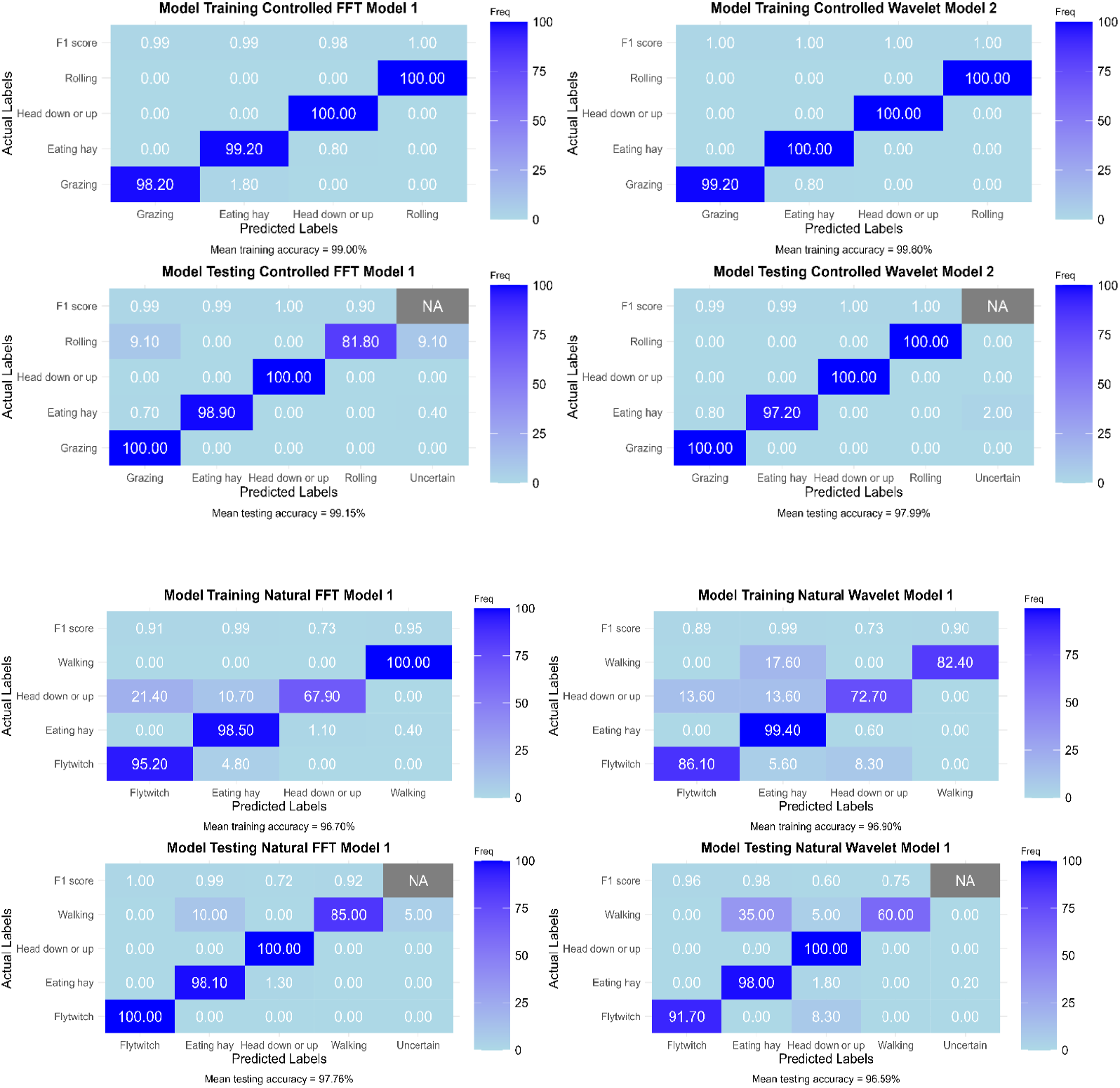
Overview of the confusion matrices pertaining to both the model training and testing of the top controlled and natural behaviour models. Accuracies are given in percentages, F1 scores are depicted as values between 0 and 1 with higher accuracies and F1 scores indicating better model performance. The colour-gradient shown on the frequence bar indicates the percentage of classification per square, with darker colours indicating a larger number of observations being classified as the corresponding movement/behaviour. Correct classifications are located on the diagonal. Mean training and testing accuracies per model are also depicted below the matrices.

## 4 Discussion

### 4.1 General results

Our results demonstrate that a single, chest mounted accelerometer (Nordic Thingy:53) coupled with an accessible machine learning (ML) platform (Edge Impulse) can classify equine gaits and natural behaviours with >95% accuracy. This is a highly significant finding given that this validates a low cost, low code input pipeline that can match the performance of more complex systems (Serra Bragança *et al*., 2020). Mean training and testing accuracy of the top models exceeded 95% for all experiments, with F1 scores exceeding 90% for gait determination and controlled behaviour experiments suggesting that gait determination and controlled behaviour models in particular may hold potential for real world applications. Previous studies have achieved comparable accuracy levels (e.g 99% for gaits) but often require multiple sensors placed on limbs (Eerdekens et al 2020) or complex custom algorithms (e.g. Serra Bragança *et al*., 2020). While limb mounted sensors may offer superior biomechanical precision for gait analysis, they are less practical for long term welfare monitoring in a paddock. Our study shows that a single, centrally located sensor can capture not just gait but also subtle “emotional state” indicators such as head position or fly twitching that may be missed by limb mounted sensors. Automated tracking methods have been used to recognise (semi-)natural behaviour in a range of species (Ladds et al., 2018; Nathan et al., 2012; Shepard et al., 2008), albeit only detecting relatively distinct movements. However, recently, both hardware and post processing algorithms have become sophisticated enough to enable the detection of more subtle behavioural differences such as gait (e.g. using a single accelerometer placed at the poll, Maisonpierre et al. (2019) were able to distinguish between standing, walking and grazing). Using the Edge Impulse platform we were able to further extend this to reliably differentiate between walking and static movements such as twitching to ward off pests (flytwitch) or moving the head up or down, something which other studies have found more challenging (Auer et al., 2021). Therefore, this work represents a significant advancement in the complexity of equine behaviour that can be defined using a relatively simple set up and a platform that enables non-experts to define models.

### 4.2 Technical considerations: Transformation type and data volume

Interestingly, we did not find a significant effect of transformation type (FFT or Wavelet) on either training or testing accuracy across our different experiments. However, we did find some differences in classification accuracy within the gait determination models (Figure 2), where the Wavelet model classified fewer movements as “uncertain”. Although previous research has shown that a Fast Fourier transform can be used to reliably detect gait (walk, trot) in horses (Roepstorff et al., 2021), the frequency information obtained from a this type of transformation is global, while a Wavelet transform offers a more localized frequency representation, which should be more effective for pattern recognition (Quchani et al., 2008). Therefore, we expected that wavelet transformation based descriptors might be superior to FFT transform based descriptors in detecting subtle changes in equine gaits. Instead, the FFT model performed comparably, and in some cases (e.g. classifying controlled behaviours) outperformed the wavelet model, when the amount of training data was reduced. This suggests that FFT transforms may handle reduced training data better than Wavelet transforms, particularly when categorising the behavioural movements captured in our controlled experiment. Rather, the limiting factor was consistently data quantity and subject diversity. For example, the reduction in data quantity characterising subtle behavioural movements, for example head up / down (model 3, Figure 4) resulted in much lower training and testing performance, highlighting that subtle behaviours require larger, more diverse datasets to prevent overfitting (Ying, 2019).

Subject variability also had a significant impact on model accuracy and F1-scores; when a specific horse was shifted to the test set, accuracy dropped for subtle movements. This confirms that individual movement signatures exist, and robust, universal models will require training across a wider range of breeds and builds than the specific Western horses used here. The effects of individual movement differences were more severe in the gait determination and controlled behaviour experiments than the natural behaviour experiments, suggesting that a broader movement signature is necessary if models are designed to detect subtle changes in gait based movement.

### 4.3 Implications for equine welfare

The ability to remotely monitor time budgets has profound implications for equine welfare assessments. Horses in sub-optimal conditions often display altered locomotion patterns such as increased restless walking, which may signal agitation (Benhajali et al., 2008; Benhajali et al., 2009), while changes in activity indexes have also been related to pain, discomfort or used to detect parturition onset (Hartmann et al., 2018). Our gait models demonstrate the potential for the automated detection of these activity shifts using a low input device without the need for labour intensive direct observation.

Crucially, our system detected semi-natural behaviours such as rolling (100% accuracy, figure 5) which can be an early indicator of disease, with a single centrally placed accelerometer rather than forelimb mounted accelerometers (e.g. see Eerdekens et al., 2024). The central placement also allowed the assessment of additional movements linked to foraging behaviour. Deviations in rolling frequency can be an early indicator of colic or other discomfort (Eerdekens et al., 2024), while reduced foraging time can be a direct indicator of negative emotional states and physiological stress (Yarnell et al., 2015). By automating the detection of these key behaviours linked to physiological and emotional states, this technology offers a scalable tool to monitor the “emotional state” of the horse (Webster 2016), with the potential to revolutionise welfare improvements in horses (Christensen et al., 2022).

### 4.4 Future directions and limitations

Our study was limited in sample size, both in terms of the number of horses used and the amount of data collected per horse and per behaviour. This is a common feature of these types of studies (e.g n = 7 in Serra Bragança et al. (2020), n = 6 in Brighton et al. (2015), n = 6 in Eerdekens et al. (2020), n = 8 in Eerdekens et al. (2024), n = 6 in Maisonpierre et al. (2019), n = 8 in Weinert et al. (2020)), yet the drop in accuracy on novel horses highlights the need for larger scale validation.

Future research should focus on determining how many horses are needed to make a truly breed agnostic model by sampling more horses across different breeds for longer periods. Expanding the range of behaviours included in the models to detect lameness (Feuser et al., 2022), indicators of distress such as excessive pawing or lying down (Eerdekens et al., 2024) or stereotypic behaviours (e.g. crib biting, weaving) would be hugely beneficial for equine welfare monitoring. Given the precision of our gait models with a single sensor compared to limb mounted sensors (e.g. Crecan et al., 2022), lameness detection would be a promising next step.

## 5 Conclusion

This study validates the Nordic Thingy:53 as a viable, accessible tool for equine behavioural analysis. We achieved >95% accuracy for gaits and natural behaviours using a single chest mounted sensor and a user-friendly machine learning platform. While data volume and training remains critical for model robustness, this approach democratises access to advanced monitoring tools. By enabling the automated detection of welfare critical behaviours, e.g. walking, rolling or positive foraging behaviour, this technology paves the way for scalable objective welfare assessment in the field.

6

## Acknowledgements

We would like to extend our sincere appreciation to the owners and caretakers who allowed data collection to be performed on their horses. Their flexibility and assistance greatly facilitated the data collection for this study. Secondly, we would like to thank the owners of the Flaming Star Ranch for providing access to indoor arenas and pastures.

## References

Aoun, C. C., Atigui, M., Brahmi, M., Gherairi, E., & Hammadi, M. (2024). Time budgets and 24 h temporal patterns variation of activities in stabled dairy dromedary camels. Applied Animal Behaviour Science, 275, 106295. 10.1016/j.applanim.2024.106295

Auer, U., Kelemen, Z., Engl, V., & Jenner, F. (2021). Activity time budgets—a potential tool to monitor equine welfare? Animals, 11(3), 850. https://www.mdpi.com/2076-2615/11/3/850

Benhajali, H., Hausberger, M., & Richard-Yris, M. A. (2007). Behavioural repertoire: its expression according to environmental conditions. In M. Hausberger, E. Søndergaard, & W. Martin-Rosset (Eds.), Horse behaviour and welfare (pp. 123–138). Wageningen Academic Publishers

Benhajali, H., Richard-Yris, M.-A., Leroux, M., Ezzaouia, M., Charfi, F., & Hausberger, M. (2008). A note on the time budget and social behaviour of densely housed horses: A case study in Arab breeding mares. Applied Animal Behaviour Science, 112(1), 196–200. 10.1016/j.applanim.2007.08.007

Benhajali, H., Richard-Yris, M. A., Ezzaouia, M., Charfi, F., & Hausberger, M. (2009). Foraging opportunity: a crucial criterion for horse welfare? Animal, 3(9), 1308–1312. 10.1017/S1751731109004820

Brighton, C., Olsen, E., & Pfau, T. (2015). Is a standalone inertial measurement unit accurate and precise enough for quantification of movement symmetry in the horse? Computer Methods in Biomechanics and Biomedical Engineering, 18(5), 527–532. 10.1080/10255842.2013.819857

Christensen, J. W., Strøm, C. G., Nicová, K., de Gaillard, C. L., Sandøe, P., & Skovgård, H. (2022). Insect-repelling behaviour in horses in relation to insect prevalence and access to shelters. Applied Animal Behaviour Science, 247, 105560. 10.1016/j.applanim.2022.105560

Crecan, C. M., Morar, I. A., Lupsan, A. F., Repciuc, C. C., Rus, M. A., & Pestean, C. P. (2022). Development of a novel approach for detection of equine lameness based on inertial sensors: A preliminary study. Sensors, 22(18), 7082.

Eerdekens, A., Deruyck, M., Fontaine, J., Martens, L., Poorter, E. D., & Joseph, W. (2020). Automatic equine activity detection by convolutional neural networks using accelerometer data. Computers and Electronics in Agriculture, 168, 105139. 10.1016/j.compag.2019.105139

Eerdekens, A., Papas, M., Damiaans, B., Martens, L., Govaere, J., Joseph, W., & Deruyck, M. (2024). Automatic early detection of induced colic in horses using accelerometer devices. Equine Veterinary Journal, n/a(n/a). 10.1111/evj.14069

Ezanno, P., Picault, S., Beaunée, G., Bailly, X., Muñoz, F., Duboz, R., Monod, H., & Guégan, J. (2021). Research perspectives on animal health in the era of artificial intelligence. Veterinary Research, 52(1), 40. 10.1186/s13567-021-00902-4

Feuser, A.-K., Gesell-May, S., Müller, T., & May, A. (2022). Artificial intelligence for lameness detection in horses—A preliminary study. Animals, 12(20), 2804.

Flannigan, G., & Stookey, J. M. (2002). Day-time time budgets of pregnant mares housed in tie stalls: a comparison of draft versus light mares. Applied Animal Behaviour Science, 78(2), 125–143. 10.1016/S0168-1591(02)00085-0

Hartmann, C., Lidauer, L., Aurich, J., Aurich, C., & Nagel, C. (2018). Detection of the time of foaling by accelerometer technique in horses (Equus caballus)-a pilot study. Reproduction in Domestic Animals, 53(6), 1279–1286. 10.1111/rda.13250

Hewson, C. J. (2003). How might veterinarians do more for animal welfare? Canadian Veterinary Journal, 44(12), 1000–1004.

Hockenhull, J., & Whay, H. R. (2014). A review of approaches to assessing equine welfare. Equine Veterinary Education, 26(3), 159–166. 10.1111/eve.12129

Huettner, T., Dollhaeupl, S., Simon, R., Baumgartner, K., & von Fersen, L. (2021). Activity Budget Comparisons Using Long-Term Observations of a Group of Bottlenose Dolphins (Tursiops truncatus) under Human Care: Implications for Animal Welfare. Animals (Basel), 11(7). 10.3390/ani11072107

Khillare, R., & Kaushal, M. (2021). Animal welfare and its importance. Agricultural Letters, 2(11), 2582–6522. https://doi.org/

Ladds, M. A., Salton, M., Hocking, D. P., McIntosh, R. R., Thompson, A. P., Slip, D. J., & Harcourt, R. G. (2018). Using accelerometers to develop time-energy budgets of wild fur seals from captive surrogates. PeerJ, 6, e5814. 10.7717/peerj.5814

Maisonpierre, I. N., Sutton, M. A., Harris, P., Menzies-Gow, N., Weller, R., & Pfau, T. (2019). Accelerometer activity tracking in horses and the effect of pasture management on time budget. Equine Veterinary Journal, 51(6), 840–845. 10.1111/evj.13130

Mench, J. A., & Mason, G. J. (1997). Behaviour. In M.C. Appleby & B. O. Hughes (Eds.), Animal Welfare (pp. 127–141). CABI Publishing.

Nathan, R., Spiegel, O., Fortmann-Roe, S., Harel, R., Wikelski, M., & Getz, W. M. (2012). Using tri-axial acceleration data to identify behavioral modes of free-ranging animals: general concepts and tools illustrated for griffon vultures. Journal of Experimental Biology, 215(6), 986–996. 10.1242/jeb.058602

Quchani, S., Moravejian, R., & mohammad kazemi, F. (2008). Gait recognition using wavelet transform Fifth international conference on information technology: new generations Las Vegas, NV, USA.

Roepstorff, C., Dittmann, M. T., Arpagaus, S., Serra Bragança, F. M., Hardeman, A., Persson-Sjödin, E., Roepstorff, L., Gmel, A. I., & Weishaupt, M. A. (2021). Reliable and clinically applicable gait event classification using upper body motion in walking and trotting horses. Journal of Biomechanics, 114, 110146. 10.1016/j.jbiomech.2020.110146

Serra Bragança, F. M., Broomé, S., Rhodin, M., Björnsdóttir, S., Gunnarsson, V., Voskamp, J. P., Persson-Sjodin, E., Back, W., Lindgren, G., Novoa-Bravo, M., Gmel, A. I., Roepstorff, C., van der Zwaag, B. J., Van Weeren, P. R., & Hernlund, E. (2020). Improving gait classification in horses by using inertial measurement unit (IMU) generated data and machine learning. Scientific Reports, 10(1), 17785. 10.1038/s41598-020-73215-9

Shepard, E., Wilson, R., Quintana, F., Laich, A., Liebsch, N., Albareda, D., Halsey, L., Gleiss, A., Morgan, D. T., & Myers, A. (2008). Identification of animal movement patterns using tri-axial accelerometry. Endangered Species Research, 10, 47–60. 10.3354/esr00084

Sweeney, C., Ennis, E., Mulvenna, M., Bond, R., & O ‘ Neill, S. (2022). How machine learning classification accuracy changes in a happiness dataset with different demographic groups. Computers, 11(5), 83. 10.3390/computers11050083

Tadich, T., Weber, C., & Nicol, C. (2013). Prevalence and factors associated with abnormal behaviors in Chilean racehorses: a direct observational study. Journal of Equine Veterinary Science, 33(2), 95–100. 10.1016/j.jevs.2012.05.059

Webster, J. (2016). Animal welfare: freedoms, dominions and “a life worth living”. Animals, 6(6), 35. https://www.mdpi.com/2076-2615/6/6/35

Weinert, J. R., Werner, J., & Williams, C. A. (2020). Validation and implementation of an automated chew sensor-based remote monitoring device as tool for equine grazing research. Journal of Equine Veterinary Science, 88, 102971. 10.1016/j.jevs.2020.102971

Whay, H. R., Main, D. C. J., Greent, L. E., & Webster, A. (2003). Animal-based measures for the assessment of welfare state of diary cattle, pigs and laying hens: Consensus of expert opinion. Animal Welfare, 12(2), 205–217.

Yarnell, K., Hall, C., Royle, C., & Walker, S. L. (2015). Domesticated horses differ in their behavioural and physiological responses to isolated and group housing. Physiology & Behavior, 143, 51–57. 10.1016/j.physbeh.2015.02.040

Ying, X. (2019). An overview of overfitting and its solutions. Journal of Physics: Conference Series, 1168(2), 022022. 10.1088/1742-6596/1168/2/022022

Zhang, L., Guo, W., Lv, C., Guo, M., Yang, M., Fu, Q., & Liu, X. (2023). Advancements in artificial intelligence technology for improving animal welfare: Current applications and research progress. Animal Research and One Health, 2. 10.1002/aro2.44

Zupan, M., Štuhec, I., & Jordan, D. (2020). The effect of an irregular feeding schedule on equine behavior. Journal of Applied Animal Welfare Science, 23(2), 156–163. 10.1080/10888705.2019.1663734

