## Supplementary data for "Validating the Nordic Thingy:53 as an Accessible Tool for Equine Behavioural Analysis using Edge AI"

Figure S1

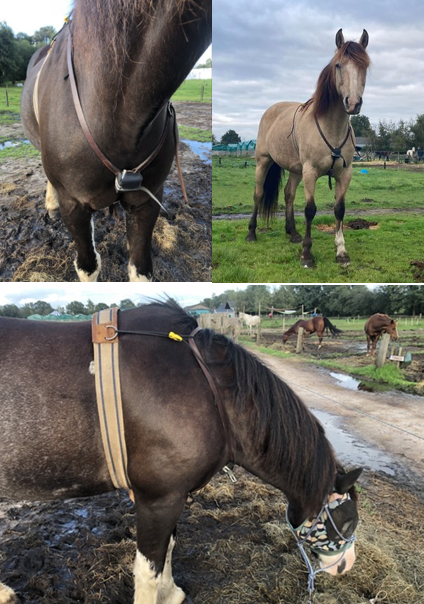

Figure S1: Placement of the Thingy:53 on the chest of the horse. The harness was designed to be minimally disturbing for the horses, while still providing a stable placement for the Thingy:53 when in motion. The Thingy:53 is located in the black leather pouch with the front of the Thingy:53 facing away from the horse’s chest and the red access port of the Thingy:53 facing towards the head of the horse, to ensure easier access to the on/off switch

Table S1

Table S1: Total amount of lunging data (in seconds) collected using the accelerometer function of the Thingy:53, which samples at a frequency of 20Hz. Amount of data per horse and per gait is provided. For the walk, trot and canter gait, the lunging direction is also provided.

| **Horse ID** | **20Hz accelerometer data** | | | | | | |
| --- | --- | --- | --- | --- | --- | --- | --- |
|  | *Idle* | *Walk L* | *Walk R* | *Trot L* | *Trot R* | *Canter L* | *Canter R* |
| *1* | 336 | 271 | 280 | 280 | 277 | 258 | 196 |
| *2* | 176 | 307 | 240 | 240 | 274 | 234 | 200 |
| *3* | 209 | 280 | 240 | 240 | 240 | 220 | 240 |
| *4* | 380 | 320 | 290 | 260 | 257 | 260 | 260 |
| *7* | / | 60 | 50 | 50 | 40 | 40 | 30 |
| *Total (sec)* | 1045 | 1518 | 1100 | 1070 | 1098 | 1012 | 926 |
| ***# horses from which data were collected*** | 4 | 5 | 5 | 5 | 5 | 5 | 5 |

Figure S2

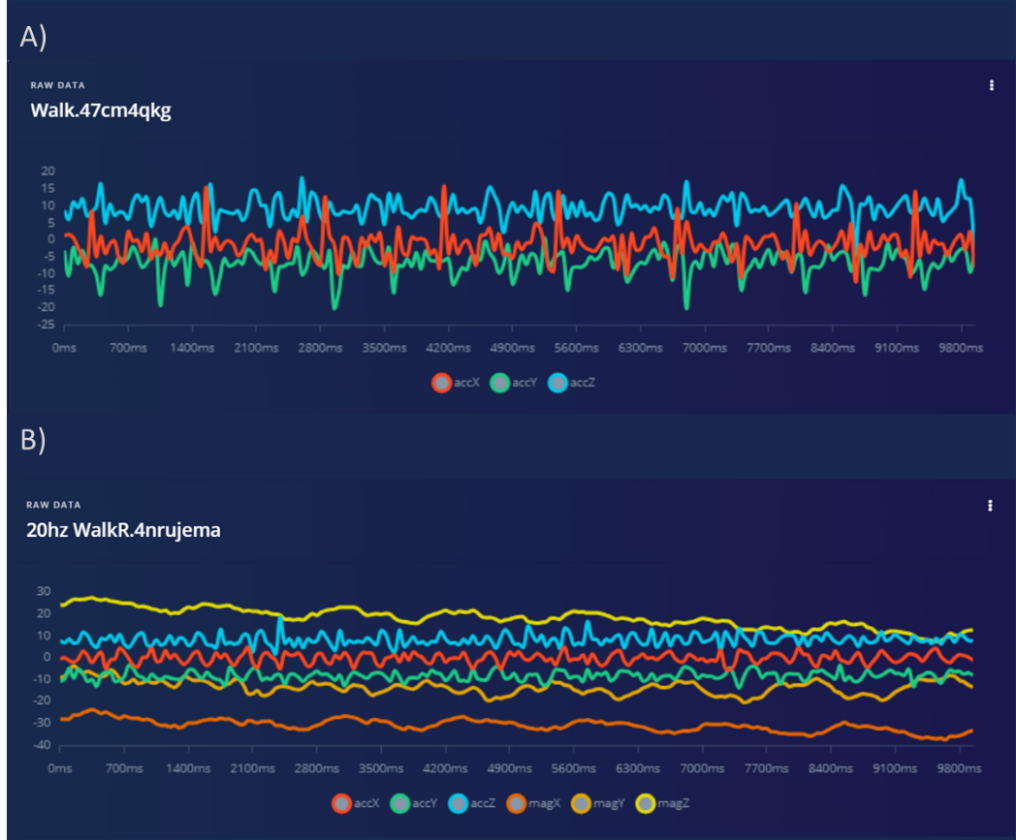

Figure S2: Example of a ten second “walk” sample collected during lunging. Data was provided by the triaxial accelerometer function of the Thingy:53 sampling at a frequency of 20Hz.

Table S2

Table S2: Overview of the amount of triaxial accelerometer data (in seconds) collected in the field with a sampling frequency of 20Hz used for the classification of controlled movements/behaviours. The amount of data collected (in seconds) is provided per horse and per movement/behaviour.

| **Controlled movements/behaviours** | **Horse ID** | | | | | **Total (sec)** | **# horses from which data were collected** |
| --- | --- | --- | --- | --- | --- | --- | --- |
|  | *1* | *2* | *3* | *4* | *16* |  |  |
| *Head down* | 22 | 32 | 30 | 106 | / | 190 | 4 |
| *Head up* | 20 | 20 | 22 | 90 | / | 152 | 4 |
| *Move head to left side* | / | 22 | / | 42 | / | 64 | 2 |
| *Move head to right side* | / | 26 | / | 42 | / | 68 | 2 |
| *Left front leg forward* | 36 | 18 | / | 2 | / | 56 | 3 |
| *Right front leg forward* | 38 | 24 | / | 2 | / | 64 | 3 |
| *Walking backwards* | 35 | 35 | / | / | / | 70 | 2 |
| *Eating hay* | 500 | 440 | 310 | 240 | / | 1450 | 4 |
| *Eating hay from haynet* | / | / | / | 320 | / | 320 | 1 |
| *Grazing* | 480 | 120 | / | 400 | / | 1000 | 3 |
| *Rolling* | 103 | 9 | / | 92 | 19 | 223 | 4 |
| Total amount of data collected per horse (sec) | 1234 | 746 | 362 | 1336 | 19 |  |  |

Figure S3

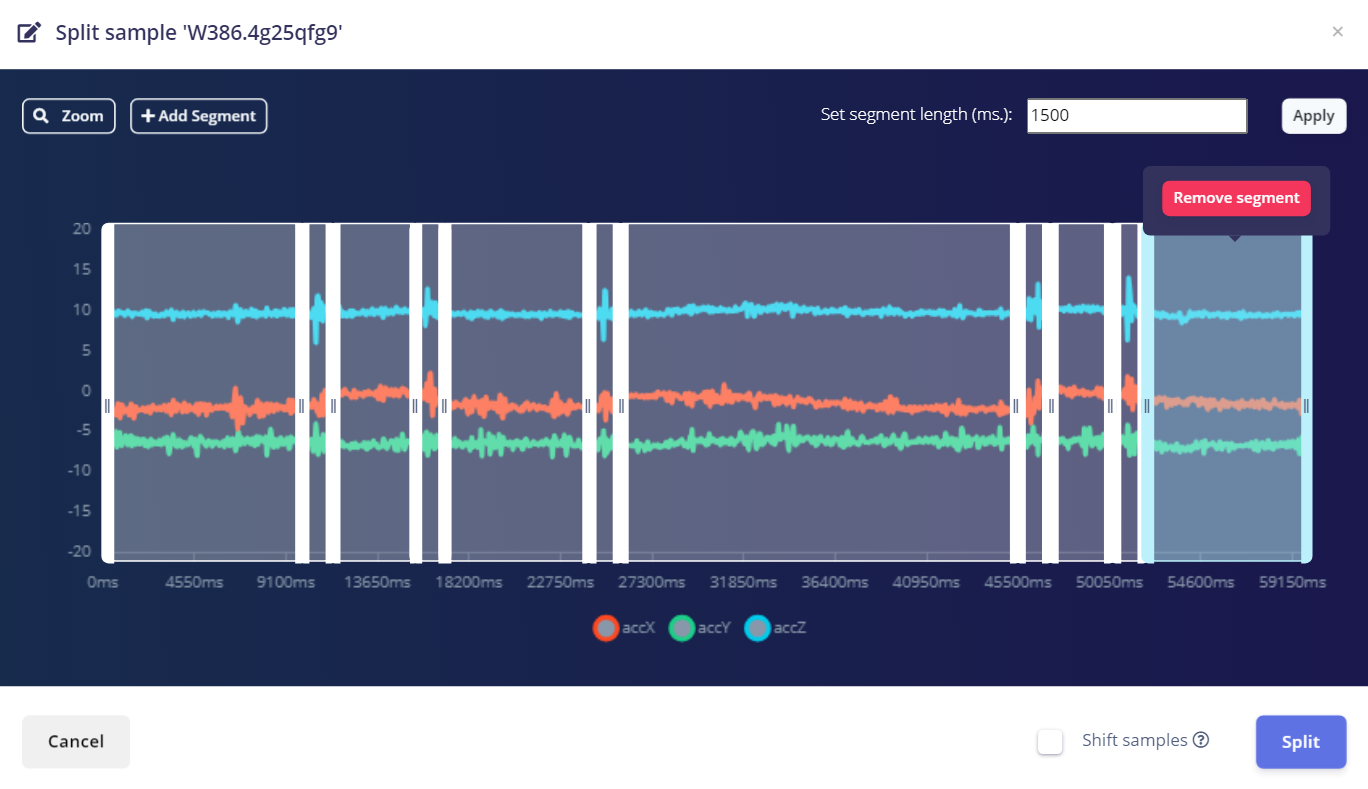

Figure S3: Splitting of a one minute sample into different movements/behaviours using the ‘split sample’ function in Edge Impulse. This specific sample would be cut into segments of grazing behaviour (segment 1,3,5,7,9 and 11) and segments in which one of the front legs is placed forward while grazing (segment 2,4,6,8 and 10).

Table S3

Table S3: Overview of the 27 natural movements/behaviours observed in the field and of the amount of triaxial accelerometer data (in seconds) collected with a sampling frequency of 20Hz used for the classification of these movements/behaviours.

| **Natural movement/behaviours** | **Abbreviation** | **Horse ID** | | | **Total (sec)** | **# horses from which data were collected** |
| --- | --- | --- | --- | --- | --- | --- |
|  |  | *1* | *2* | *5* |  |  |
| *Placing the left front leg forward* | *Llf* | 109 | 64 | 15 | 188 | 3 |
| *Placing the right front leg forward* | *Rlf* | 122 | 53 | 12 | 187 | 3 |
| *Placing the left front leg back* | *Llb* | 4 | 1 | 4 | 9 | 3 |
| *Placing the right front leg back* | *Rlb* | 12 | 6 | 3 | 21 | 3 |
| *Placing the head down* | *Hd* | 68 | 31 | 22 | 121 | 3 |
| *Placing the head up* | *Hu* | 77 | 25 | 13 | 115 | 3 |
| *Swatting at flies with the left front leg* | *Llfs* | 17 | 24 | / | 41 | 2 |
| *Swatting at flies with the right front leg* | *Rlfs* | 11 | 56 | / | 67 | 2 |
| *Swatting at flies with the left hind leg* | *Lhlfs* | 22 | 6 | 3 | 31 | 3 |
| *Swatting at flies with the right hind leg* | *Rhlfs* | 14 | 6 | / | 20 | 2 |
| *Twitching due to flies* | *Ft* | 90 | 171 | 14 | 275 | 3 |
| *Scratching the head against the left front leg* | *Hsll* | 9 | 49 | / | 58 | 2 |
| *Scratching the head against the right front leg* | *Hsrl* | / | 14 | / | 14 | 1 |
| *Scratching the left flank with the head* | *Hsls* | 24 | 6 | / | 30 | 2 |
| *Scratching the right flank with the head* | *Hsrs* | 25 | 8 | / | 33 | 2 |
| *Shaking the head* | *Hs* | 40 | 98 | 27 | 165 | 3 |
| *Grazing* | */* | 638 | / | / | 638 | 1 |
| *Eating hay* | *Hay* | 2092 | 1181 | 726 | 3999 | 3 |
| *Feeding* | *Feed* | 132 | / | 133 | 265 | 2 |
| *Urinating* | */* | 9 | / | / | 9 | 1 |
| *Drinking* | */* | 16 | 43 | 15 | 74 | 3 |
| *Walking* | */* | 79 | 44 | 30 | 153 | 3 |
| *Walking backwards* | *Wb* | 16 | 19 | / | 35 | 2 |
| *Standing still* | *Idle* | 31 | 311 | / | 342 | 2 |
| *Producing a spluttering sound by exhaling through the nostrils* | *Splutter* | / | 15 | 6 | 21 | 2 |
| *Pushing hay away with the head* | *Phwh* | / | 4 | 14 | 18 | 2 |
| *Touch head to left flank* | *Htls* | / | / | 5 | 5 | 1 |
| Total amount of data collected per horse (sec) |  | 3657 | 2235 | 1042 |  |  |

Figure S4

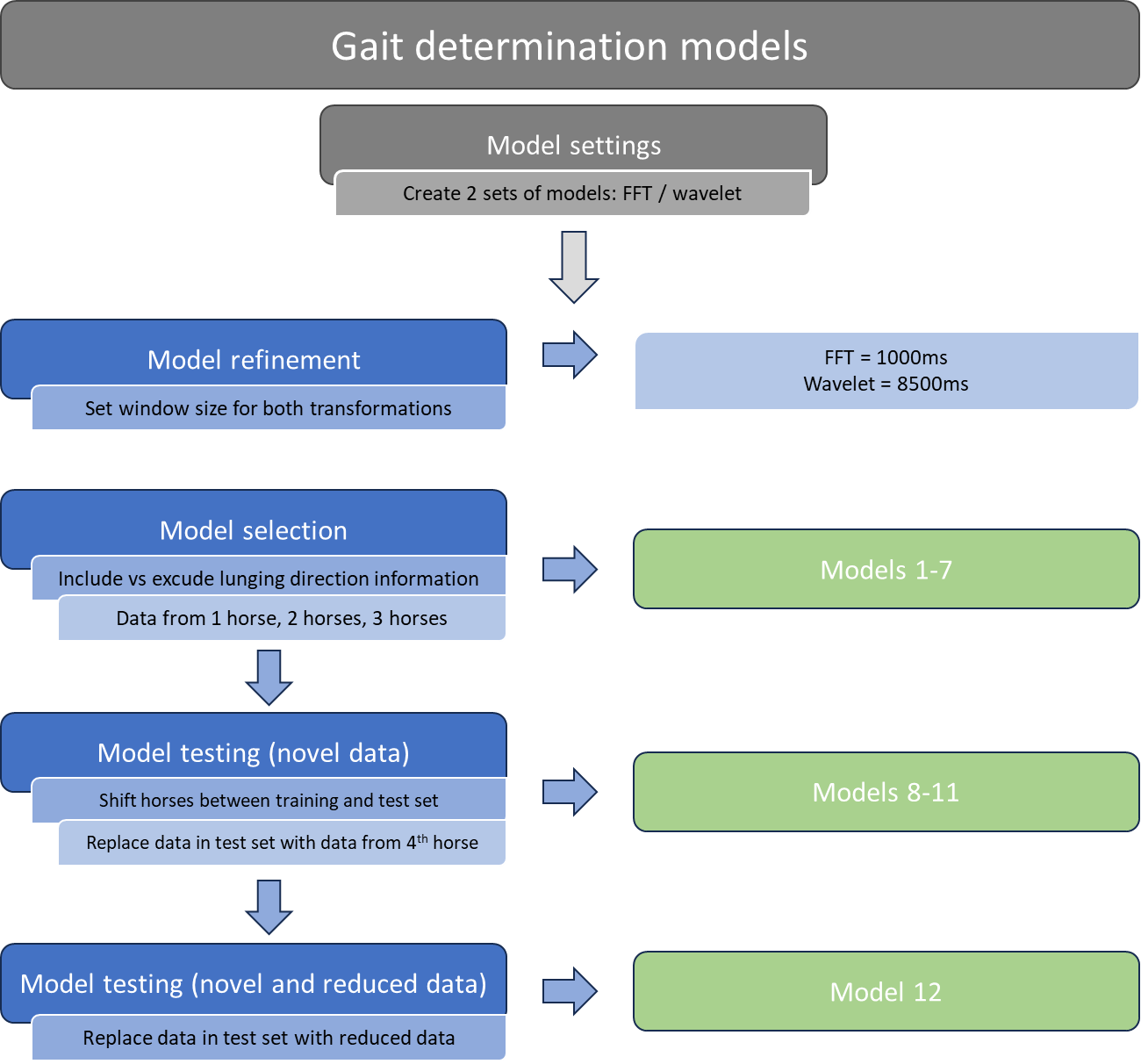

Figure S4: Flow chart detailing the steps taken during the construction of gait determination models. First, the spectral analysis type was selected being either a Fast Fourier transform or a Wavelet transform. Hereafter, a window size was chosen for both transforms during model refinement. Consequent model selection consisted of the in- and exclusion of the lunging direction information as well as increasing the amount of training data (from data obtained from 1 horse to data obtained from 2 and 3 horses). After model selection, the robustness of the gait determination models was determined by first testing the robustness of the model to novel data (models 8-11; horses shifted between training and test data sets) and then to reduced novel data (model 12; horse with reduced data placed in the test set).

Figure S5

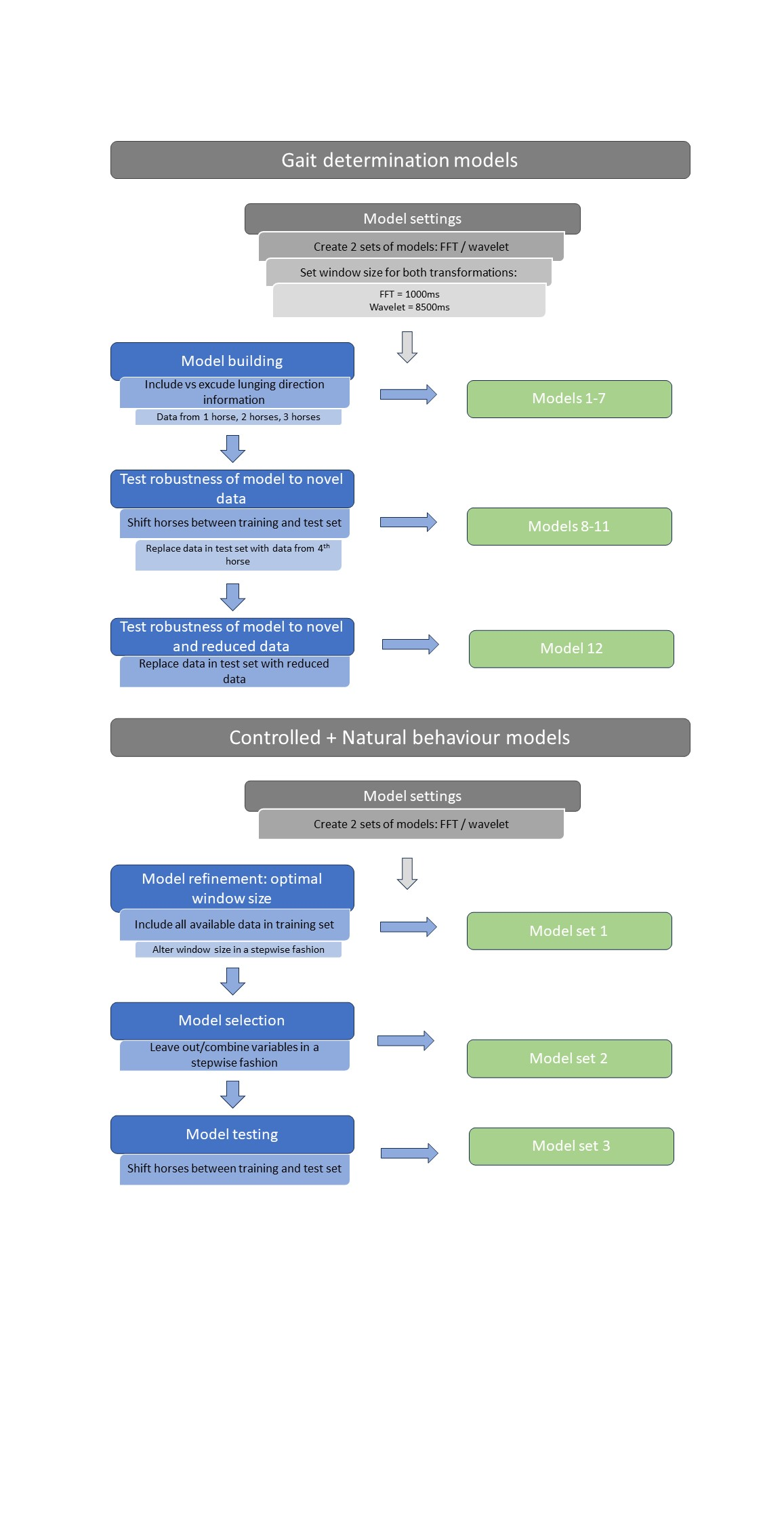

Figure S5: Flow chart detailing the steps taken during the construction of controlled and natural behaviour models. First, the spectral analysis type was selected being either a Fast Fourier transform or a Wavelet transform. Hereafter, the window size for both transforms was refined in a stepwise fashion during model refinement. Consequent model selection consisted of leaving out or combining variables in a stepwise fashion. Lastly, to test the robustness of the models to the introduction of novel data, horses were shifted between the training and test set, creating model set 3.

Table S4

| **Transformation** | **FFT** | ***Model*** | ***1*** | ***2*** | ***3*** | ***4*** | ***5*** | ***6*** | ***7*** | ***8*** | ***9*** | ***10*** | ***11*** | ***12*** | ***13*** | ***14*** | ***15*** | ***16*** | ***17*** | ***18*** | ***19*** | ***20*** | ***21*** | ***22*** |
| --- | --- | --- | --- | --- | --- | --- | --- | --- | --- | --- | --- | --- | --- | --- | --- | --- | --- | --- | --- | --- | --- | --- | --- | --- |
|  |  | ***Window size (ms)*** | 1000 | 2000 | 3000 | 4000 | 5000 | 6000 | 7000 | 8000 | 9000 | 10000 | 11000 | 12000 | 13000 | 14000 | 15000 | 16000 | 17000 | 18000 | 19000 | 20000 | 30000 | 40000 |
|  |  | ***Training accuracy (%)*** | 67.80 | 76.00 | 78.70 | 82.00 | 84.40 | 88.10 | 84.10 | 85.70 | 86.80 | 87.90 | 89.40 | 89.70 | 87.90 | 87.00 | 88.90 | 89.40 | 88.80 | 91.00 | 89.00 | 87.50 | 82.10 | 50.60 |
|  | **Wavelet** | ***Model*** | ***1*** | ***2*** | ***3*** | ***4*** | ***5*** | ***6*** | ***7*** | ***8*** | ***9*** | ***10*** | ***11*** | ***12*** | ***13*** | ***14*** | ***15*** | ***16*** | ***17*** | ***18*** | ***19*** | ***20*** | ***21*** |  |
|  |  | ***Window size (ms)*** | 3200 | 4000 | 5000 | 6000 | 7000 | 8000 | 9000 | 10000 | 11000 | 12000 | 13000 | 14000 | 14500 | 15000 | 16000 | 17000 | 18000 | 19000 | 20000 | 30000 | 40000 |  |
|  |  | ***Training accuracy (%)*** | 81.90 | 83.00 | 85.90 | 88.40 | 83.60 | 87.00 | 87.00 | 88.30 | 90.40 | 89.50 | 89.20 | 91.00 | 89.10 | 88.70 | 90.90 | 89.80 | 92.00 | 90.40 | 90.30 | 83.00 | 48.30 |  |

Table S4: Overview of the change in training accuracy imposed on the controlled behaviour models, comprising model set 1, when the window size was altered in a stepwise fashion to determine the optimal window size. For the models on which the optimal window size is based, training accuracy values are indicated in grey.

Table S5

Table S5: Overview of the change in training accuracy imposed on the natural behaviour models comprising model set 1 when the window size was altered in a stepwise fashion to determine the optimal window size. For the models on which the optimal window size is based, training accuracy values are indicated in grey.

| **Transformation** | **FFT** | ***Model*** | ***1*** | ***2*** | ***3*** | ***4*** | ***5*** | ***6*** | ***7*** | ***8*** | ***9*** | ***10*** | ***11*** | ***12*** | ***13*** | ***14*** |
| --- | --- | --- | --- | --- | --- | --- | --- | --- | --- | --- | --- | --- | --- | --- | --- | --- |
|  |  | ***Window size (ms)*** | 1000 | 2000 | 3000 | 4000 | 5000 | 6000 | 7000 | 8000 | 9000 | 10000 | 11000 | 12000 | 13000 | / |
|  |  | ***Training accuracy (%)*** | 69.80 | 78.30 | 78.80 | 80.00 | 75.60 | 77.70 | 76.30 | 74.90 | 75.50 | 75.30 | 72.70 | 75.40 | 71.00 | / |
|  | **Wavelet** | ***Model*** | ***1*** | ***2*** | ***3*** | ***4*** | ***5*** | ***6*** | ***7*** | ***8*** | ***9*** | ***10*** | ***11*** | ***12*** | ***13*** | ***14*** |
|  |  | ***Window size (ms)*** | 3200 | 3400 | 3600 | 3800 | 4000 | 5000 | 6000 | 7000 | 8000 | 9000 | 10000 | 11000 | 12000 | 13000 |
|  |  | ***Training accuracy (%)*** | 80.40 | 79.40 | 80.50 | 82.10 | 80.00 | 77.50 | 78.30 | 77.80 | 79.10 | 76.90 | 75.70 | 73.80 | 76.70 | 72.20 |

Table S6

Table S6: Overview of the data present in the controlled behaviour models, belonging to model set 2. An overview of training accuracy values obtained in each model selection step are also given. A distinction is made between models using an Fast Fourier transform and models using a Wavelet transform. Top models are indicated in grey.

| **Model** | **# Movements/behaviours contained in model** | **Alteration** | **Reasoning behind alteration** | **Training set** | **Test set** | **Training accuracy (%)** | |
| --- | --- | --- | --- | --- | --- | --- | --- |
|  |  |  |  | # Horses | | FFT | Wavelet |
| *1* | 11 | / | / | 5 | 0 | 91.00 | 92.00 |
| *2* | 10 | Combined “left front leg forward” and “right front leg forward” | - F1 score “right front leg forward” = 0 - Similar movements | 5 | 0 | 92.00 | 91.60 |
| *3* | 8 | Removed “move head to left side”+”move head to right side” | - F1 scores = 0 - Similar movements | 5 | 0 | 92.90 | 95.40 |
| *4* | 7 | Combined “head down” and “head up” | - Similar movements - FFT: “head up” = lowest F1 score - Wavelet: “front leg forward” 60% categorized as "head up” | 5 | 0 | 95.00 | 95.20 |
| *5* | 6 | Removed “eating hay from haynet” | - FFT: lowest accuracy - Wavelet: Lowest F1 score | 5 | 0 | 96.80 | 98.20 |
| *6* | 5 | Removed “front leg forward” | - Lowest F1 score | 5 | 0 | 99.70 | 99.40 |
| *7* | 4 | Removed “walking backwards | - No accuracy or F1 score obtained | 5 | 0 | 99.70 | 99.70 |

Table S7

Table S7: Overview of the data present in the natural behaviour models belonging to model set 2. An overview of training accuracy values obtained in each model selection step are also given. A distinction is made between models using an Fast Fourier transform and models using a Wavelet transform. Top models are indicated in grey.

| **Model** | **# Movements/behaviours contained in model** | **Alteration** | **Reasoning behind alteration** | **Training set** | **Test set** | **Training accuracy (%)** | |
| --- | --- | --- | --- | --- | --- | --- | --- |
|  |  |  |  | # Horses | | FFT | Wavelet |
| *1* | 27 | / | / | 3 | 0 | 80.00 | / |
| *2* | 26 | Removed htls | - Insufficient data | 3 | 0 | 79.60 | / |
| *3* | 25 | Combined hsll and hsrl | - Insufficient data for hsrl - F1 score = 0 | 3 | 0 | 80.20 | 79.30 |
| *4* | 24 | Combined llb and rlb | - Insufficient data - F1 score = 0 | 3 | 0 | 80.20 | 79.70 |
| *5* | 23 | Combined llfs and rlfs | - F1 score = 0 - Only collected for 2 horses | 3 | 0 | 79.10 | 78.30 |
| *6* | 22 | Combined lhlfs and rhlfs | - F1 score = 0 | 3 | 0 | 79.30 | 79.00 |
| *7* | 21 | Combined hsls and hsrs | - F1 score = 0 | 3 | 0 | 78.80 | 78.50 |
| *8* | 20 | Combined (llb + rlb) and wb | - F1 score = 0 | 3 | 0 | 79.50 | 79.30 |
| *9* | 19 | Removed peeing | - Insufficient data | 3 | 0 | 79.40 | 80.20 |
| *10* | 18 | Removed (llb+rlb+wb) | - Insufficient data - F1 score = 0 | 3 | 0 | 79.10 | 80.50 |
| *11* | 17 | Combined hay and phwh | - F1 score phwh = 0 - FFT: Phwh was 100% classified as hay - Insufficient data for phwh | 3 | 0 | 77.90 | 79.90 |
| *12* | 16 | Removed splutter | - Insufficient data | 3 | 0 | 80.20 | 79.20 |
| *13* | 15 | Removed drinking | - F1 score = 0 | 3 | 0 | 78.50 | 80.40 |
| *14* | 14 | Removed feed | - F1 score = 0 | 3 | 0 | 88.10 | 84.10 |
| *15* | 13 | Combined (lhlfs+rhlfs) and (llfs+rlfs) | - Similar movements - Low F1 scores | 3 | 0 | 88.10 | 84.50 |
| *16* | 12 | Removed (hsls+hsrs) | - F1 score = 0 | 3 | 0 | 86.00 | 83.80 |
| *17* | 11 | Removed (hsll+hsrl) | - F1 score = 0 | 3 | 0 | 86.10 | 87.20 |
| *18* | 10 | Combined llf and rlf | - Similar movements - Low F1 scores | 3 | 0 | 86.80 | 86.50 |
| *19* | 9 | Removed (lhlfs+rhlfs+llfs+rlfs | - Low F1 score | 3 | 0 | 88.40 | 87.70 |
| *20* | 8 | Combined hd and hu | - Lowest F1 scores | 3 | 0 | 89.00 | 88.80 |
| *21* | 7 | Removed hs | - Lowest F1 score | 3 | 0 | 89.70 | 92.90 |
| *22* | 6 | Removed (llf+rlf) | - Lowest F1 score | 3 | 0 | 93.20 | 92.20 |

Table S8

Table S8: Overview of the amount of accelerometer data (in seconds) present in controlled behaviour model 7, used for the classification of controlled movements/behaviours in model set 3.

| **Controlled movements/behaviours** | **Horse ID** | | | | | **Total (sec)** | **# horses from which data were collected** |
| --- | --- | --- | --- | --- | --- | --- | --- |
|  | *1* | *2* | *3* | *4* | *16* |  |  |
| *Head down or up* | 42 | 52 | 52 | 196 | / | 342 | 4 |
| *Eating hay* | 500 | 440 | 310 | 240 | / | 1490 | 4 |
| *Grazing* | 480 | 120 | / | 400 | / | 1000 | 3 |
| *Rolling* | 103 | 9 | / | 92 | 19 | 223 | 4 |
| Total (sec) / horse | 1125 | 621 | 362 | 928 | 19 | 3055 | / |

Table S9

Table S9: Overview of the amount of accelerometer data (in seconds) present in natural behaviour model 22, used for the classification of natural movements/behaviours in model set 3.

| **Natural movements/behaviours** | **Horse ID** | | | **Total (sec)** | **# horses from which data were collected** |
| --- | --- | --- | --- | --- | --- |
|  | *1* | *2* | *5* |  |  |
| *Head down or up* | 145 | 56 | 35 | 236 | 3 |
| *Eating hay* | 2092 | 1181 | 726 | 3999 | 3 |
| *Walking* | 79 | 44 | 30 | 153 | 3 |
| *Flytwitch* | 90 | 171 | 14 | 275 | 3 |
| Total (sec) / horse | 2406 | 1452 | 805 | 4663 | / |

Table S10

Table S10: Overview of the ID amount of data present in the gait determination models for each gait and lunging direction. Additionally, a distinction is made between the horses and data contained in the training or test set of the model. For each model, the transformation and window size used are also given, as well as information on the potential inclusion of a novel horse in the test set and lunging direction in the models.

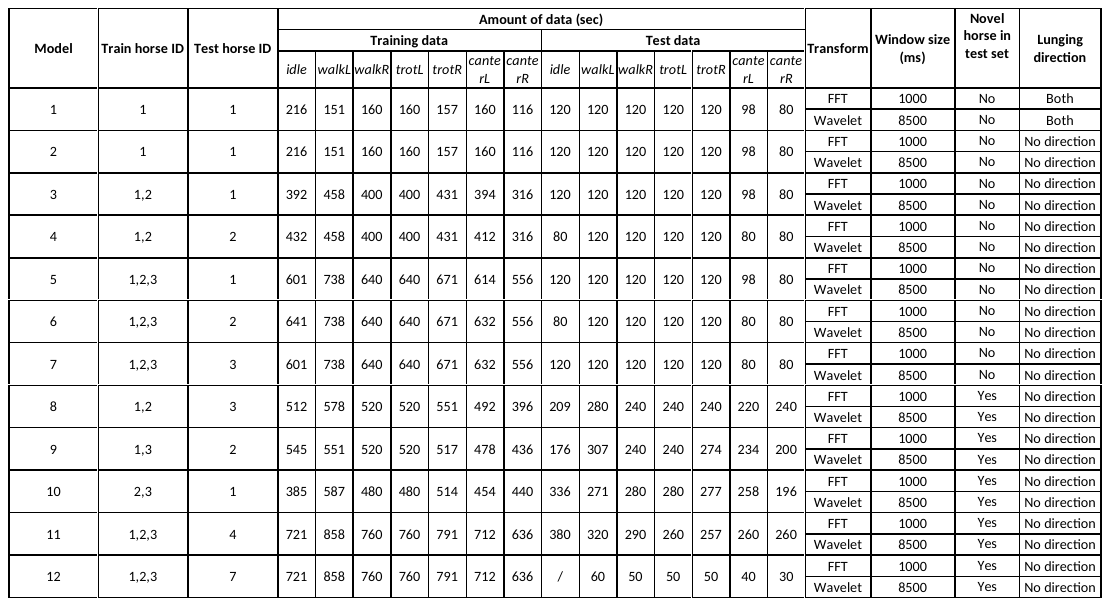

Table S11

Table S11: Overview of the structure of the models used in model set 3 of the controlled behaviour experiment (indicated in blue) and natural behaviour experiment (indicated in orange). The training accuracy and model testing accuracy values (%) of the models are also given. Accuracy values belonging to the top models are indicated in bold.

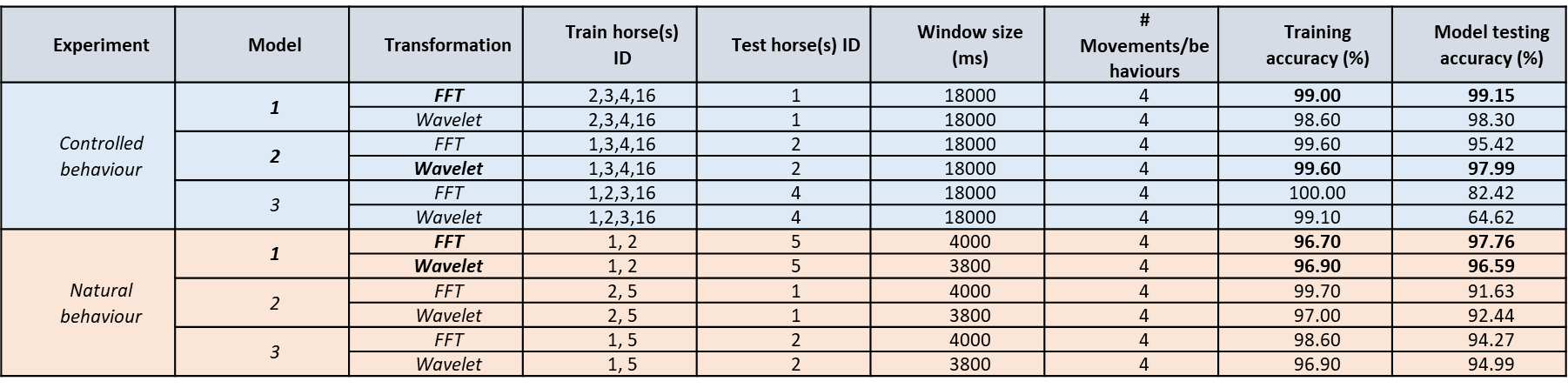

.

Figure S6

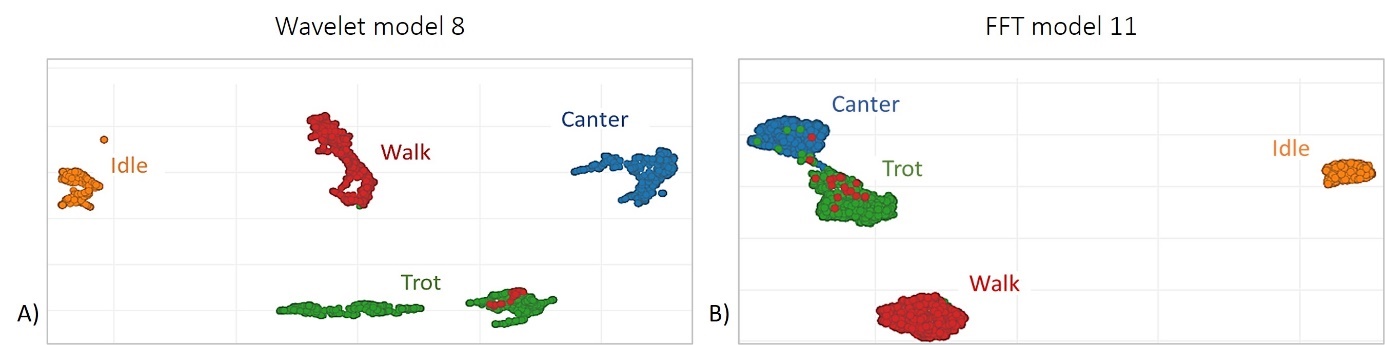

Figure S6: Feature explorer plots obtained for A) Wavelet model 8 and B) FFT model 11. Explorer plots represent a visualization of the features, i.e. the outputs of the processing block, obtained by the models. The degree of separation among the classes indicates the model’s ability to reliably differentiate between the different gaits.

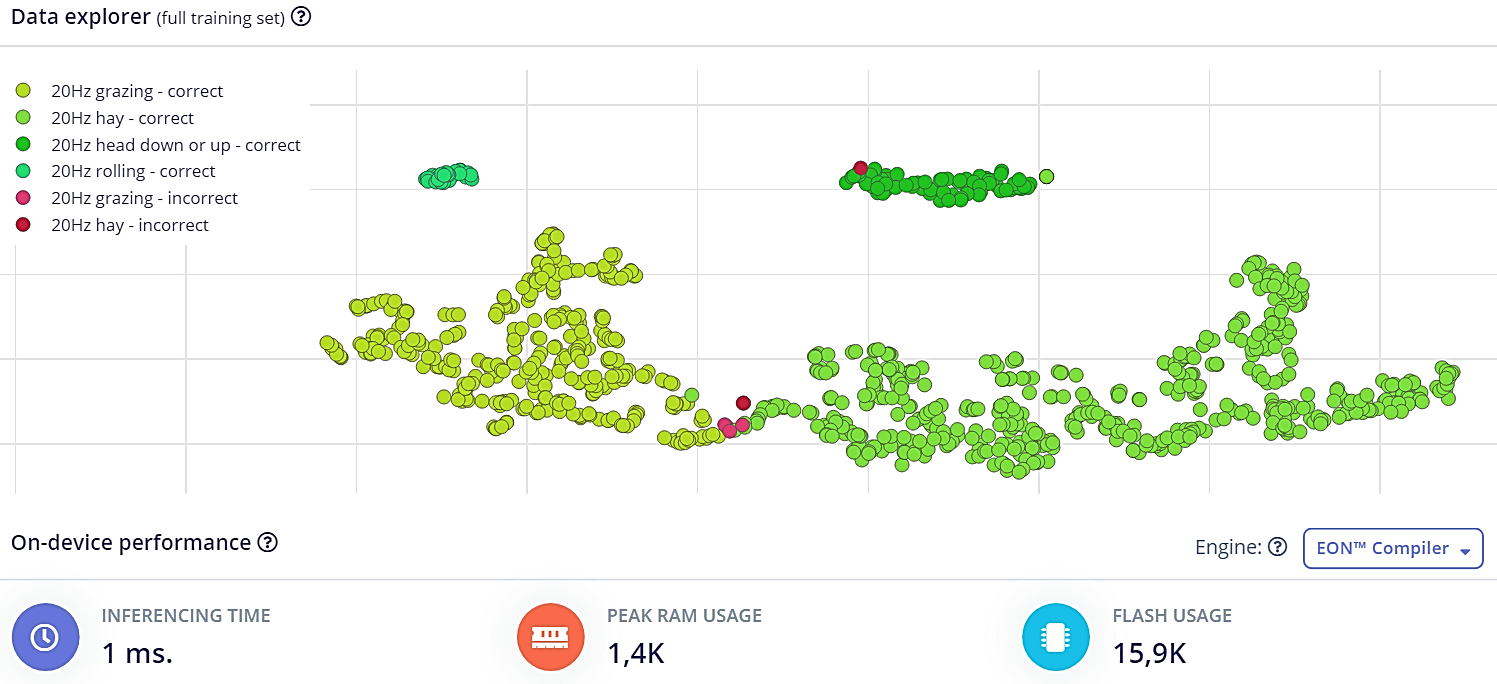

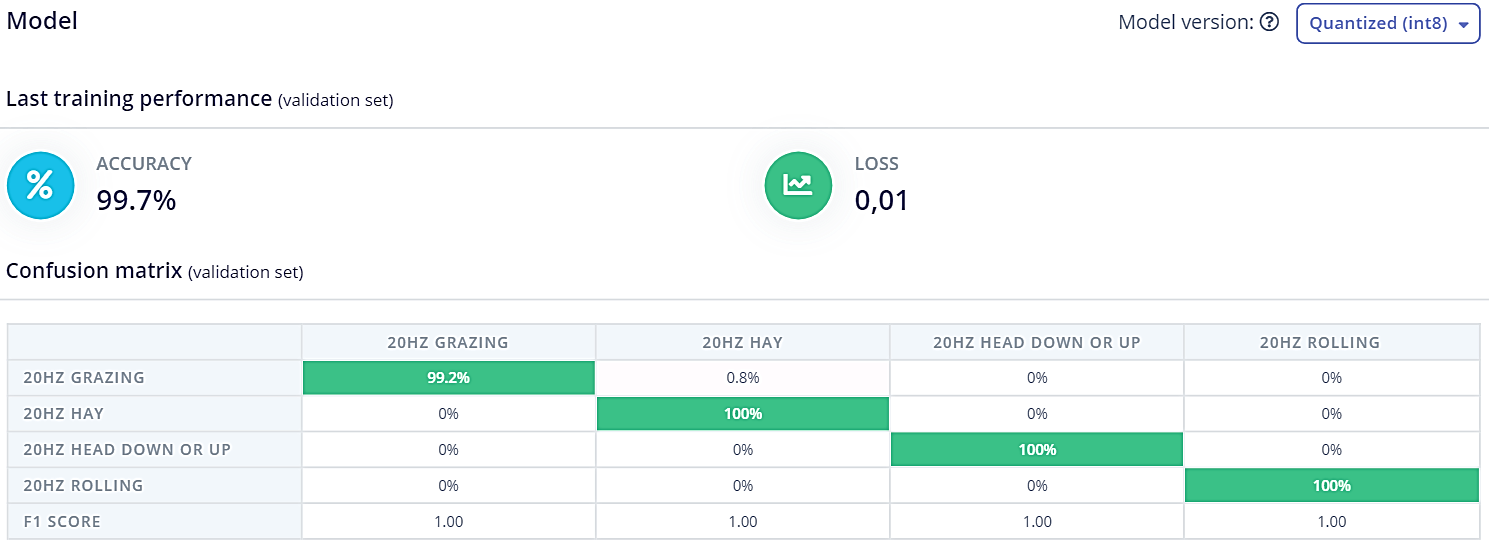
Figure S7

Figure S7: Confusion matrix obtained during model training of the final FTT model from controlled behaviour model set 2, namely model 7.

Figure S8

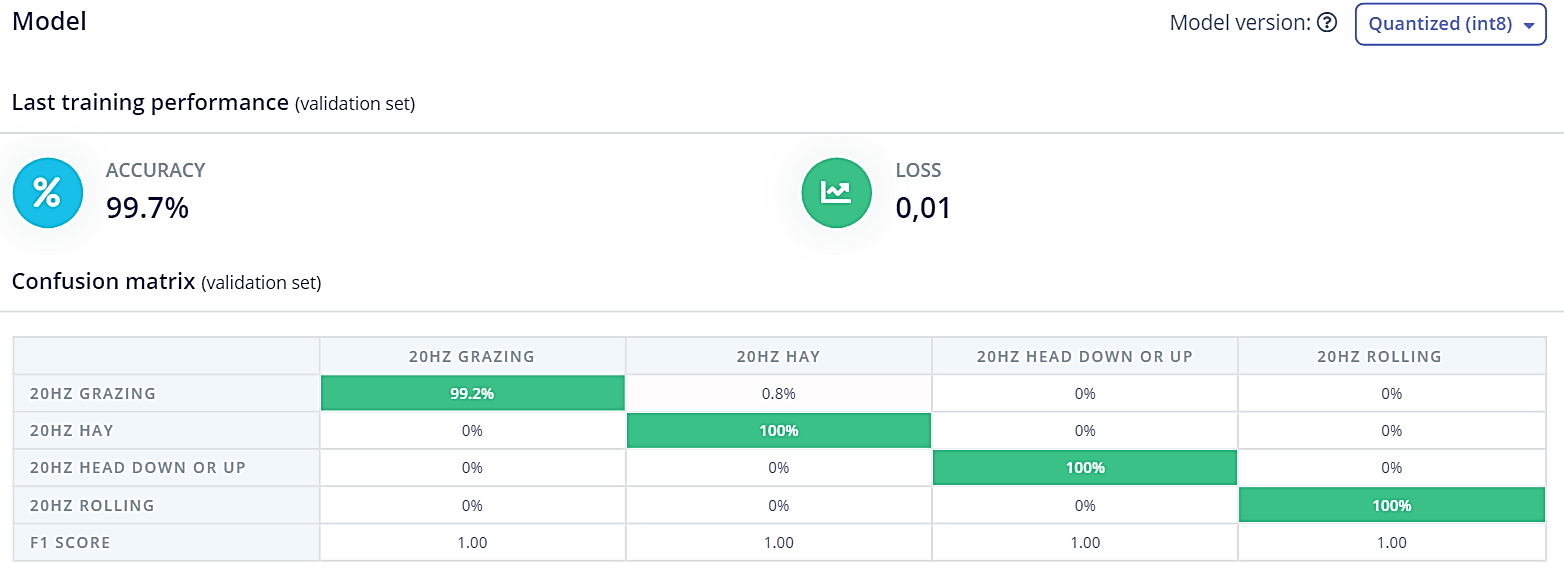

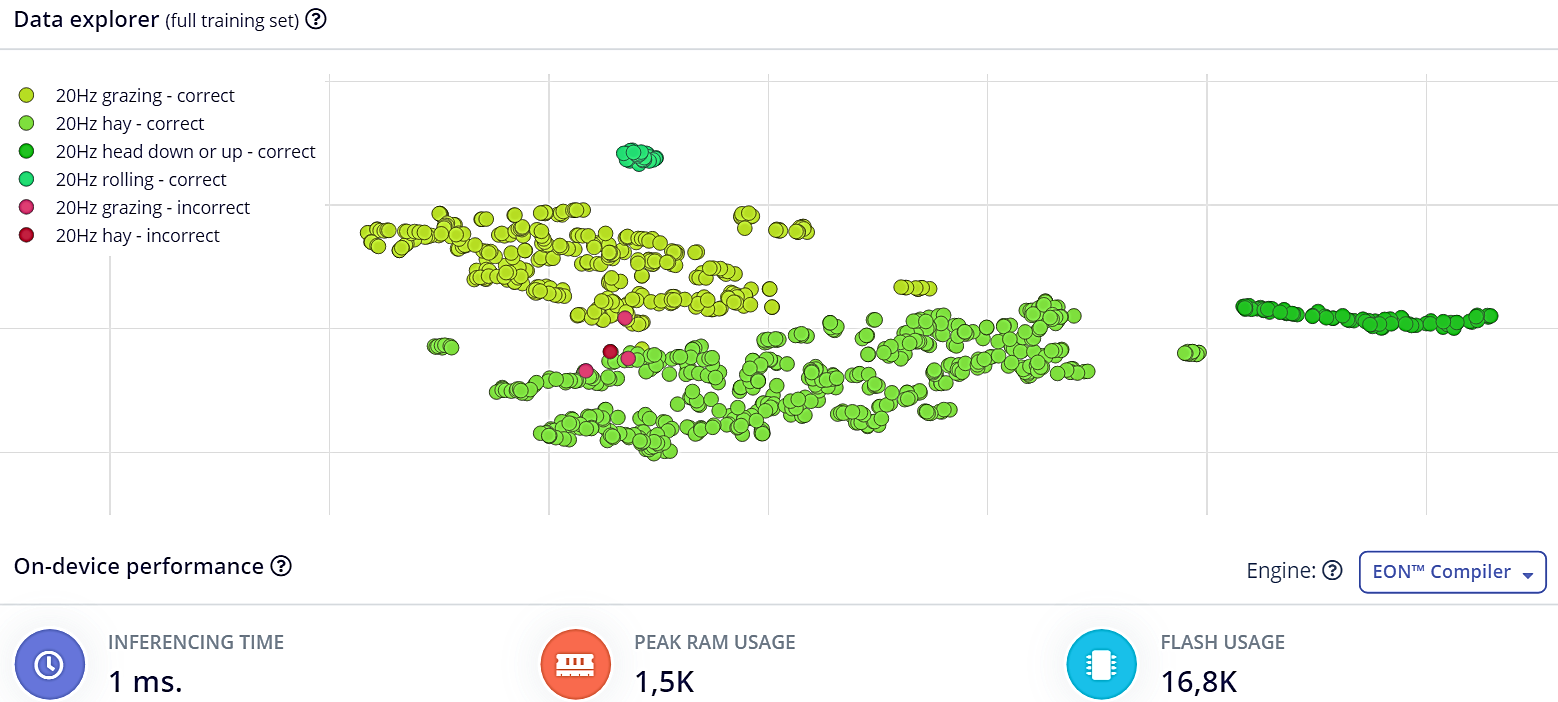

Figure S8: Confusion matrix obtained during model training of the final accelerometer wavelet model from controlled behaviour model set 2, namely model 7.

Figure S9

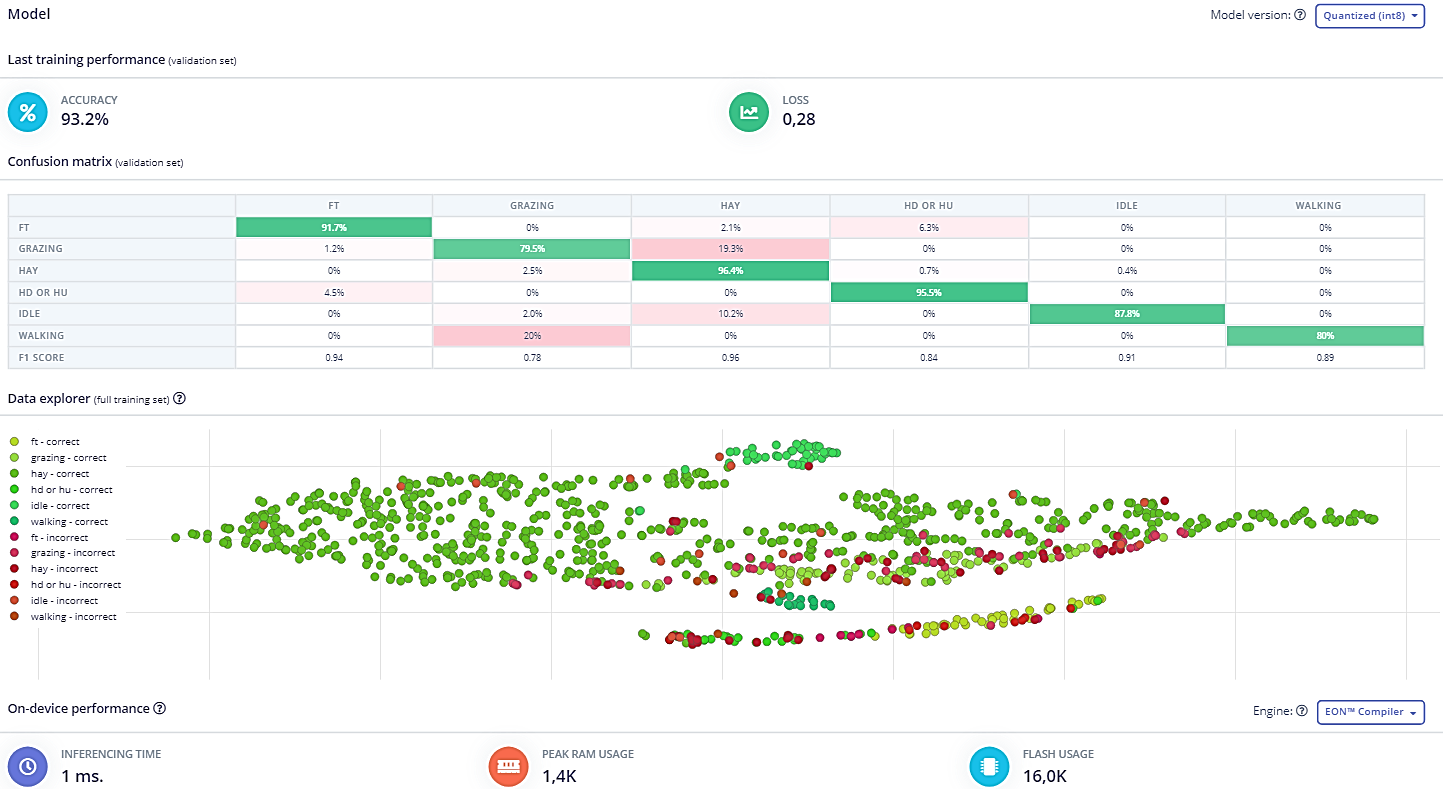

Figure S9: Confusion matrix obtained during model training of the final FFT model from natural behaviour model set 2, namely model 22.

Figure S10

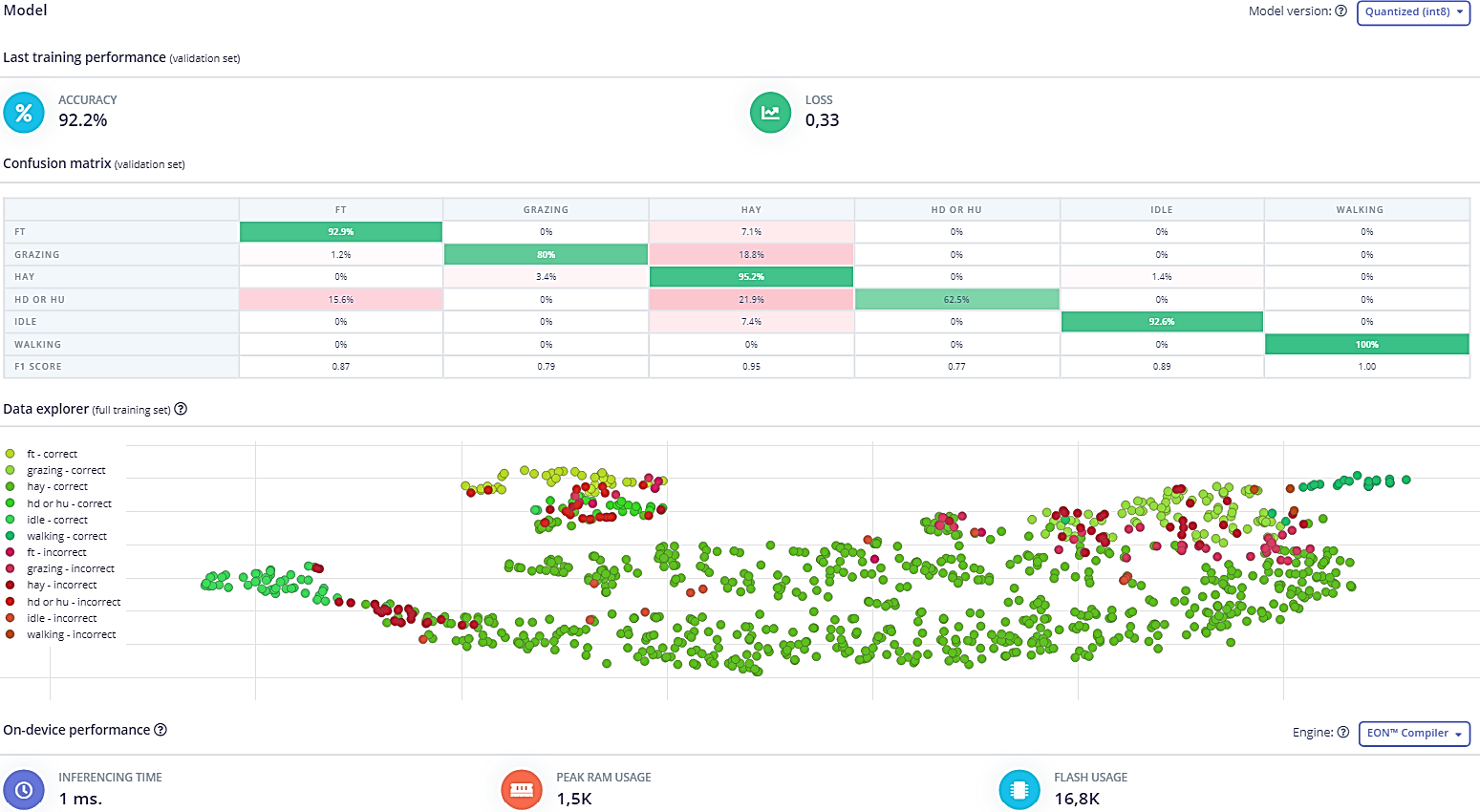

Figure S10: Confusion matrix obtained during model training of the final wavelet model from natural behaviour model set 2, namely model 22.

Figure S11

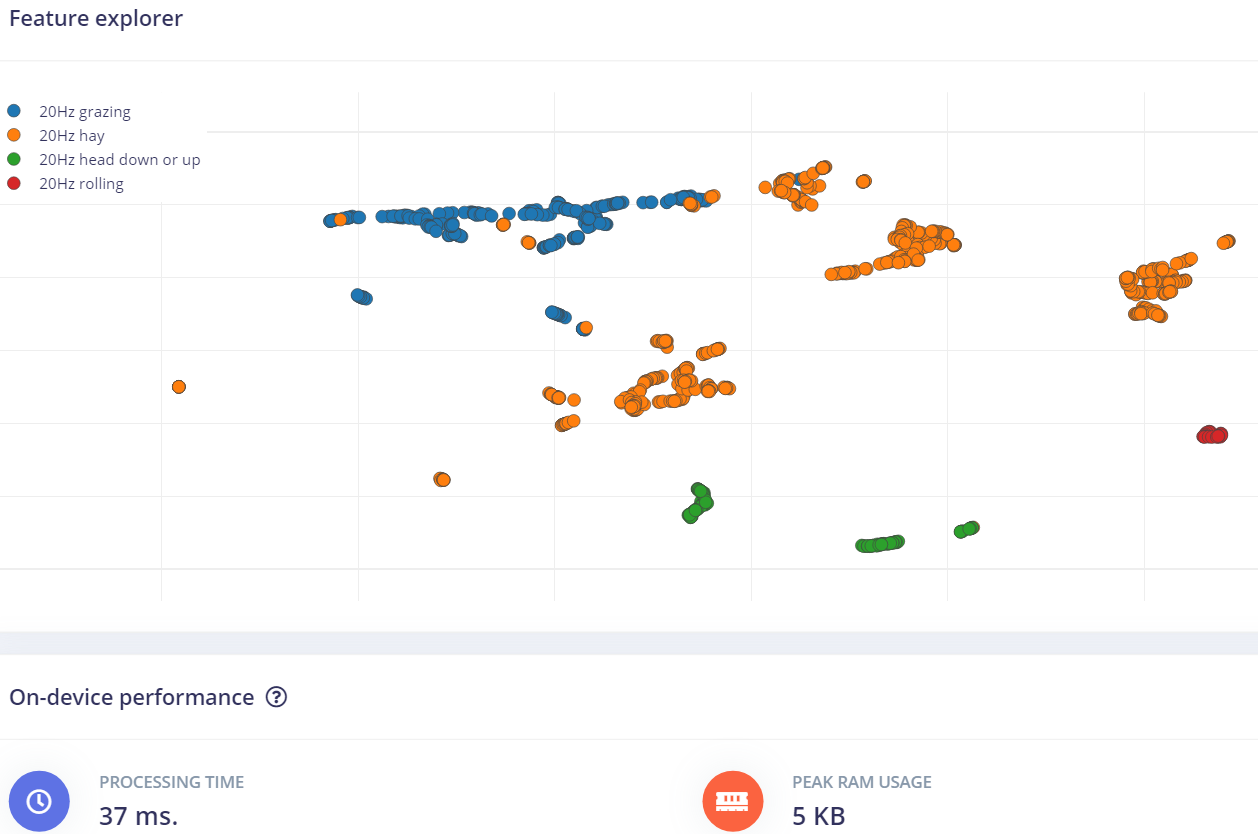

Figure 11: Feature explorer plots obtained for top FFT model 1 within the controlled behaviour experiment. Plots represent a visualization of the features, i.e. the outputs of the processing block, obtained by the models. The degree of separation among the classes indicates the model’s ability to reliably differentiate between the different controlled movements/behaviours.

OR

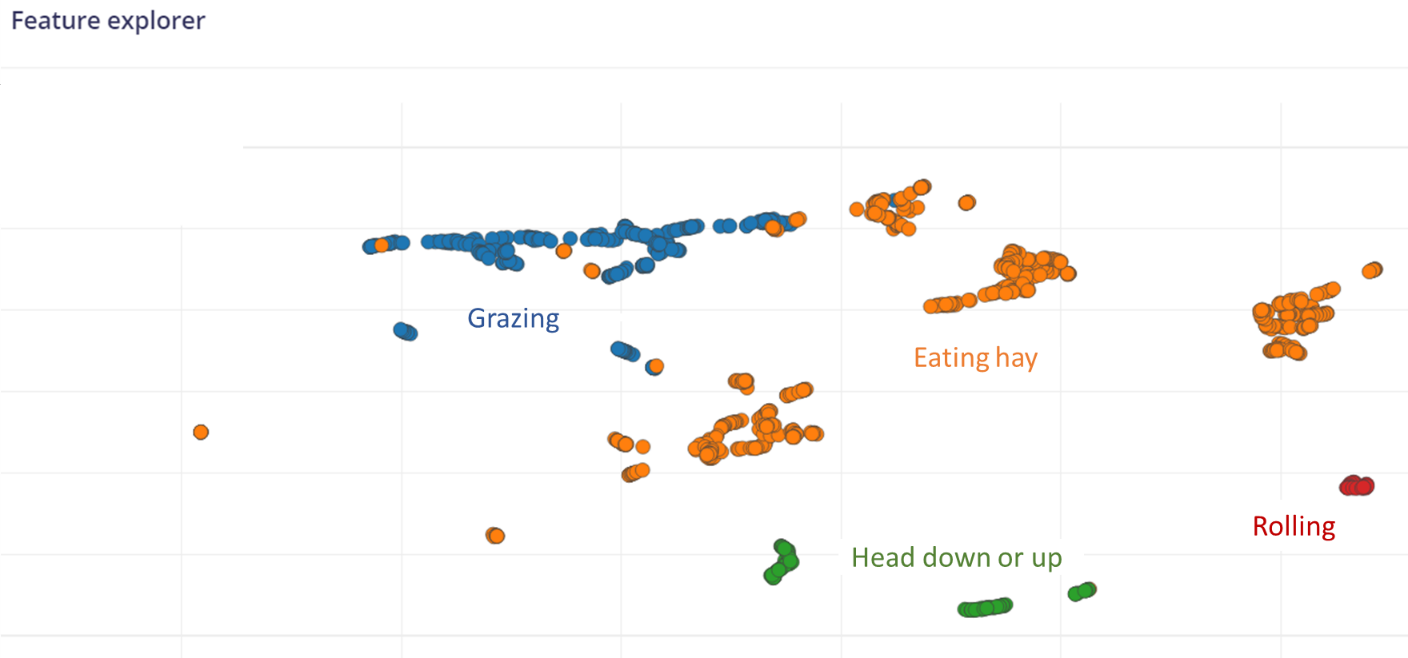

Figure S11: Feature explorer plots obtained for top FFT model 1 within the controlled behaviour experiment. Plots represent a visualization of the features, i.e. the outputs of the processing block, obtained by the models. The degree of separation among the classes indicates the model’s ability to reliably differentiate between the different controlled movements/behaviours.

Figure S12

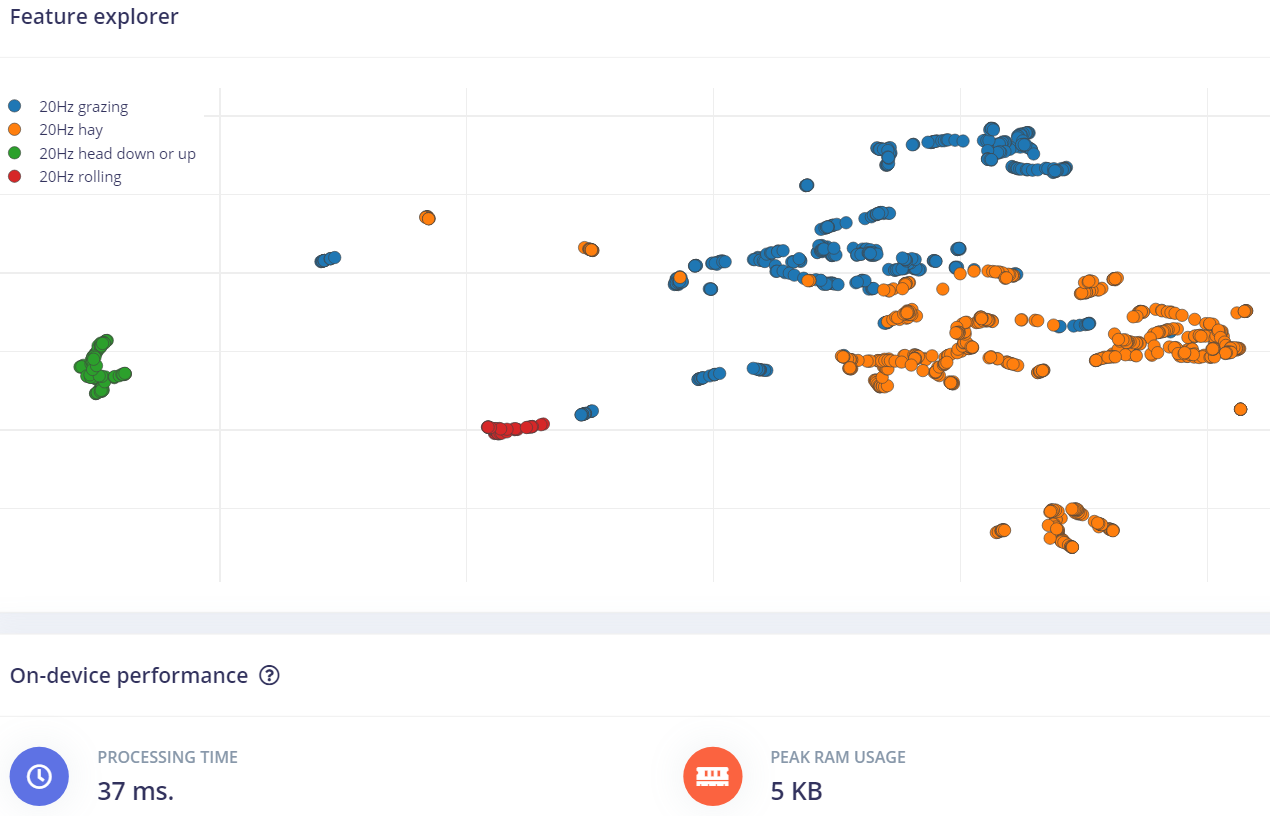

Figure S12: Feature explorer plot obtained for top Wavelet model 2 within the controlled behaviour experiment. The plot represents a visualization of the features, i.e. the outputs of the processing block, obtained by the model. The degree of separation among the classes indicates the model’s ability to reliably differentiate between the different controlled movements/behaviours.

OR

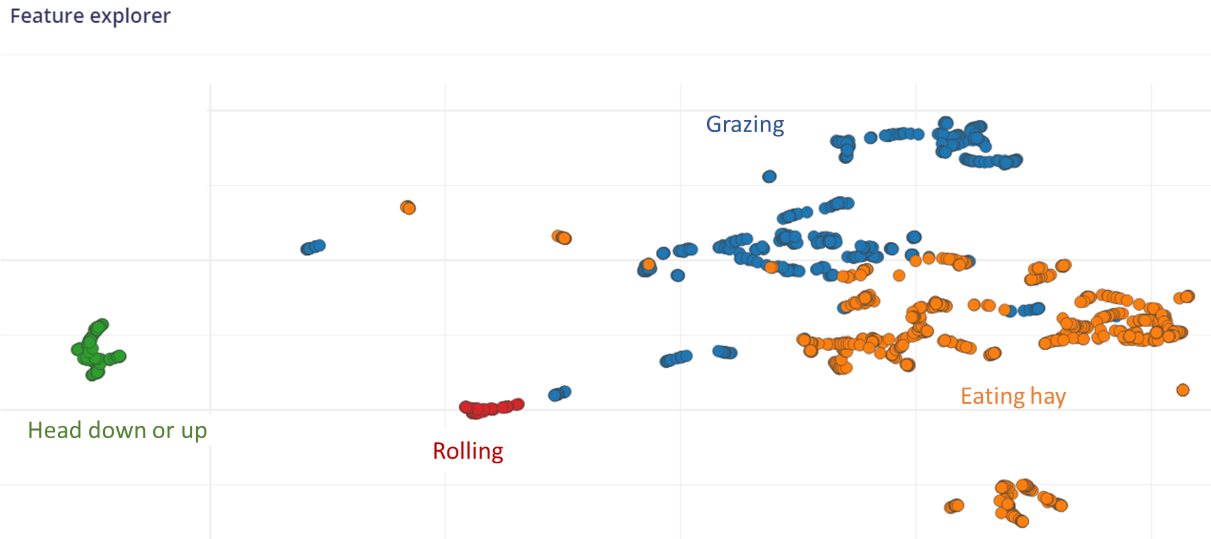

Figure S12: Feature explorer plot obtained for top Wavelet model 2 within the controlled behaviour experiment. The plot represents a visualization of the features, i.e. the outputs of the processing block, obtained by the model. The degree of separation among the classes indicates the model’s ability to reliably differentiate between the different controlled movements/behaviours.

Figure S13

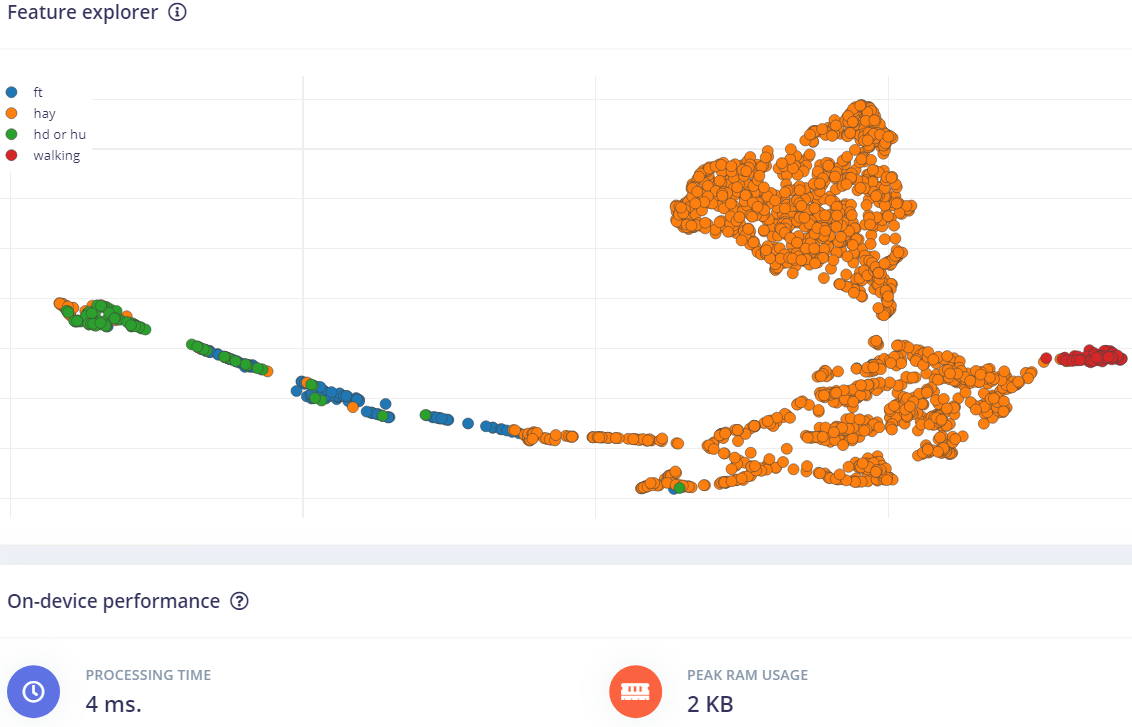

Figure S13: Feature explorer plot obtained for top FFT model 1 within the natural behaviour experiment. The plot represents a visualization of the features, i.e. the outputs of the processing block, obtained by the model. The degree of separation among the classes indicates the model’s ability to reliably differentiate between the different natural movements/behaviours.

OR

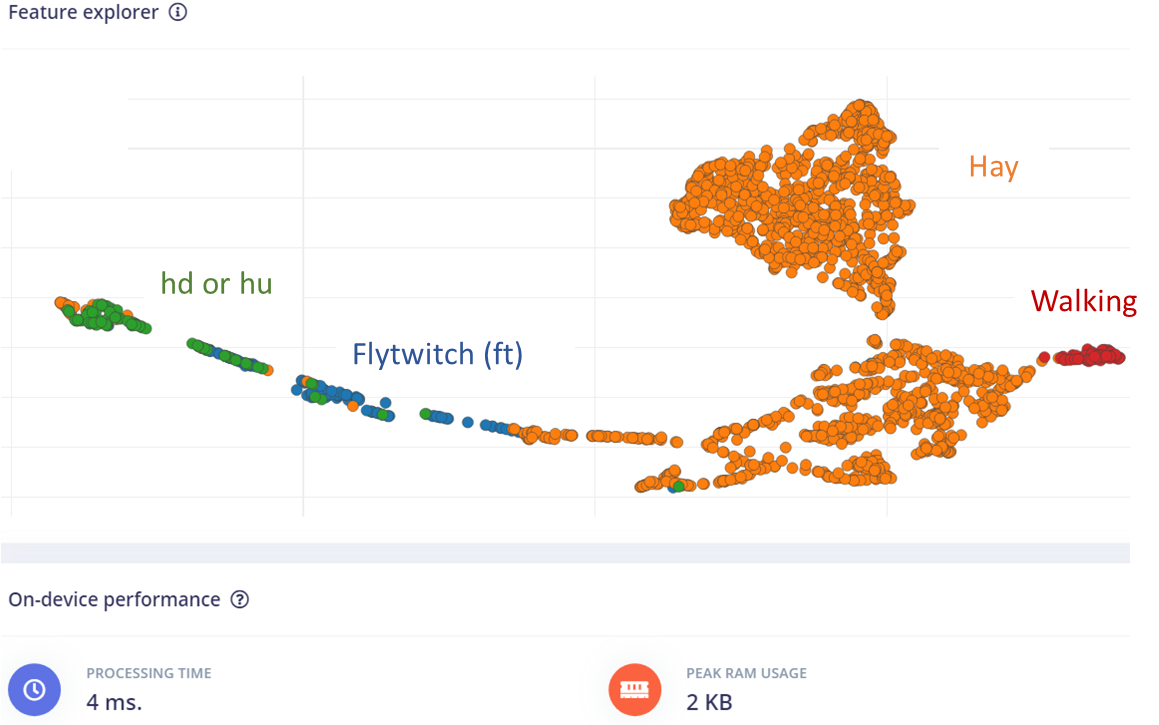

Figure S13: Feature explorer plot obtained for top FFT model 1 within the natural behaviour experiment. The plot represents a visualization of the features, i.e. the outputs of the processing block, obtained by the model. The degree of separation among the classes indicates the model’s ability to reliably differentiate between the different natural movements/behaviours.

Figure S14

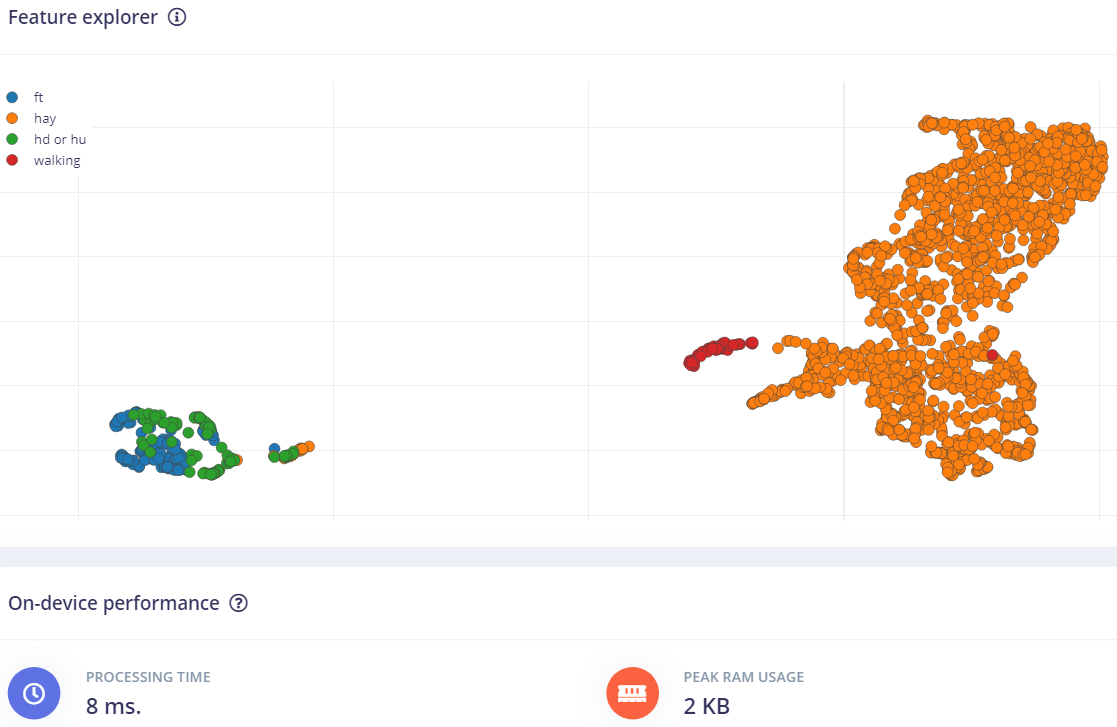

Figure S14: Feature explorer plot obtained for top Wavelet model 1 within the natural behaviour experiment. The plot represents a visualization of the features, i.e. the outputs of the processing block, obtained by the model. The degree of separation among the classes indicates the model’s ability to reliably differentiate between the different natural movements/behaviours.

OR

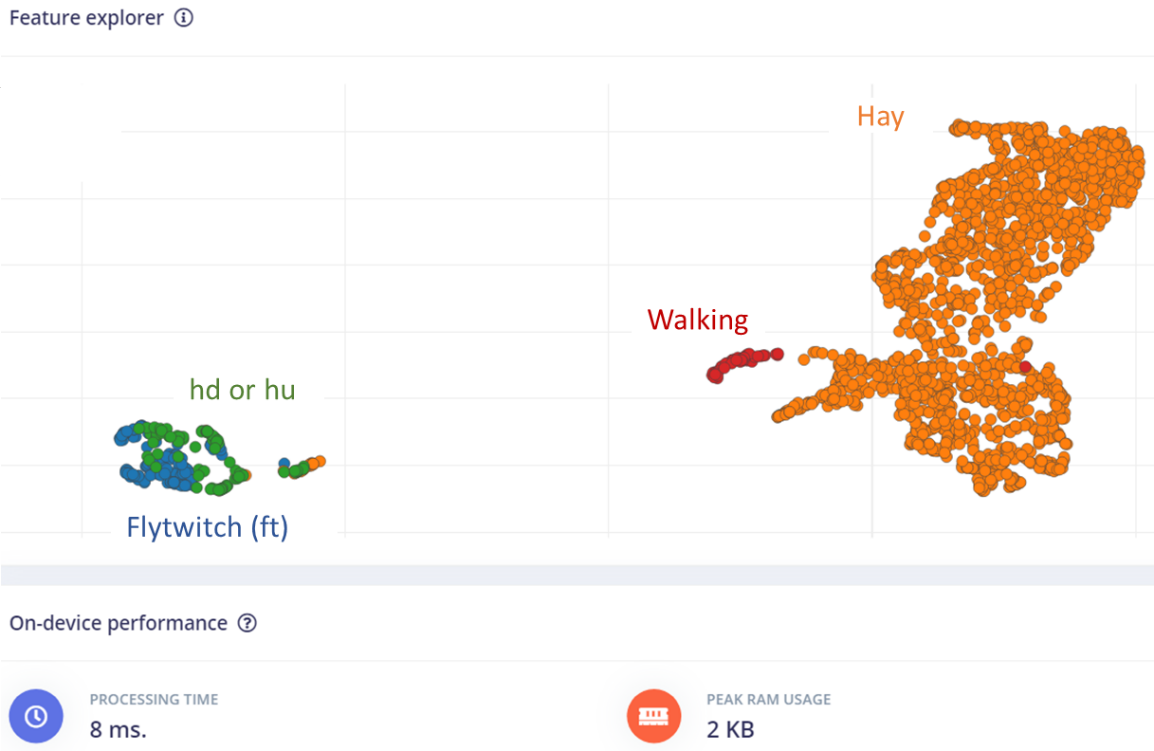

Figure S14: Feature explorer plot obtained for top Wavelet model 1 within the natural behaviour experiment. The plot represents a visualization of the features, i.e. the outputs of the processing block, obtained by the model. The degree of separation among the classes indicates the model’s ability to reliably differentiate between the different natural movements/behaviours.
